# A biodegradable “one-for-all” nanoparticle for multimodality imaging and enhanced photothermal treatment of breast cancer

**DOI:** 10.1101/2023.11.28.568885

**Authors:** Jessica C. Hsu, Diego Barragan, Alexander E. Tward, Maryam Hajfathalian, Ahmad Amirshaghaghi, Katherine J. Mossburg, Derick N. Rosario-Berríos, Mathilde Bouché, Alexander K. Andrianov, E. James Delikatny, David P. Cormode

## Abstract

Silver sulfide nanoparticles (Ag_2_S-NP) have been proposed for various optical-based biomedical applications, such as near-infrared fluorescence (NIRF) imaging, photoacoustics (PA) and photothermal therapy (PTT). However, their absorbance is relatively low in the NIR window used in these applications, and previous formulations were synthesized using toxic precursors under harsh conditions and have clearance issues due to their large size. Herein, we synthesized sub-5 nm Ag_2_S-NP and encapsulated them in biodegradable, polymeric nanoparticles (AgPCPP). All syntheses were conducted using biocompatible reagents in the aqueous phase and under ambient conditions. We found that the encapsulation of Ag_2_S-NP in polymeric nanospheres greatly increases their NIR absorbance, resulting in enhanced optical imaging and photothermal heating effects. We therefore found that AgPCPP have potent contrast properties for PA and NIRF imaging, as well as for computed tomography (CT). We demonstrated the applicability of AgPCPP nanoparticles as a multimodal imaging probe that readily improves the conspicuity of breast tumors *in vivo*. PTT was performed using AgPCPP with NIR laser irradiation, which led to significant reduction in breast tumor growth and prolonged survival compared to free Ag_2_S-NP. Lastly, we observed a gradual decrease in AgPCPP retention in tissues over time with no signs of acute toxicity, thus providing strong evidence of safety and biodegradability. Therefore, AgPCPP may serve as a “one-for-all” theranostic agent that degrades into small components for excretion once the diagnostic and therapeutic tasks are fulfilled, thus providing good prospects for translation to clinical use.

**TOC graphic:** 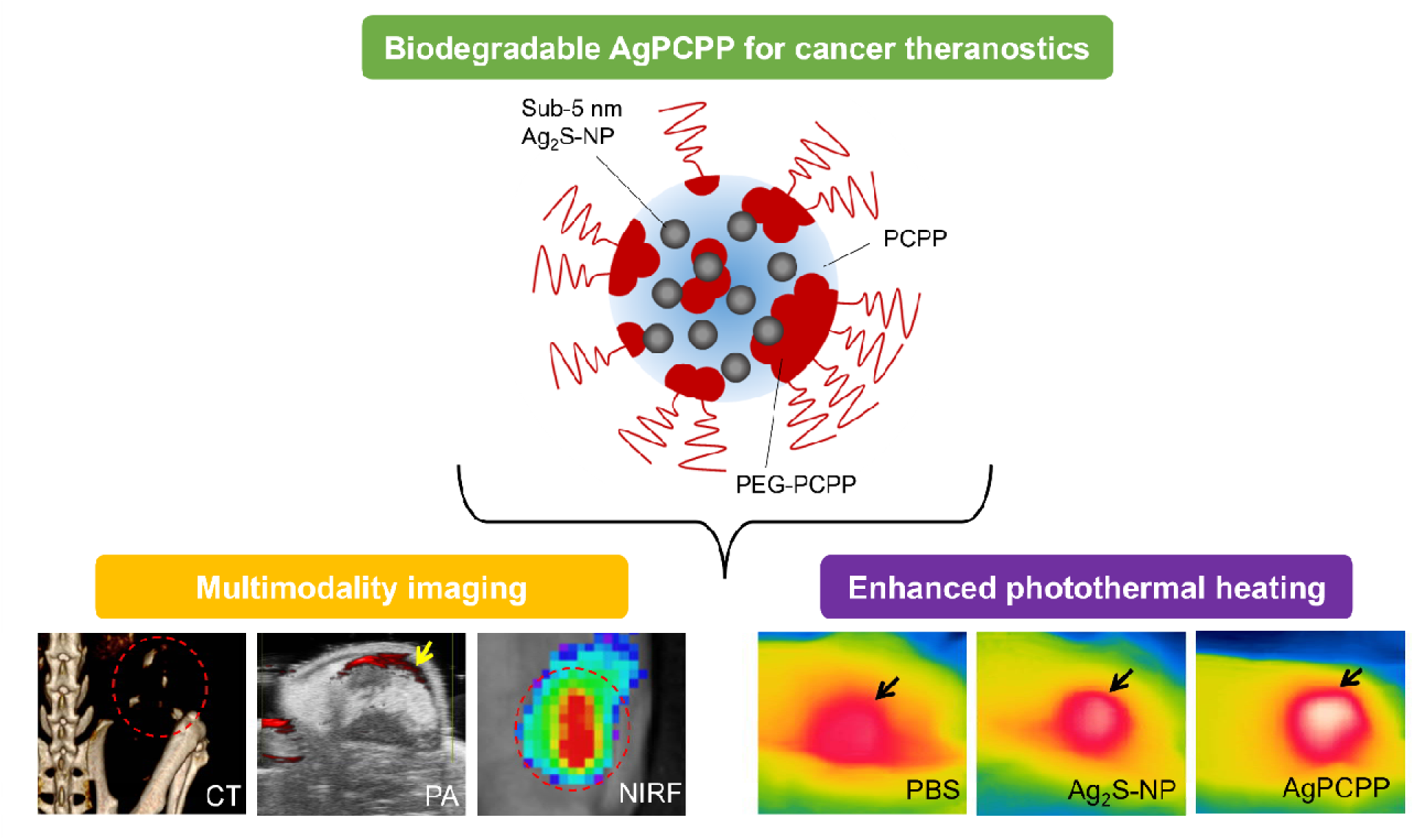

---

Nano-theranostics are typically composed of both diagnostic and therapeutic agents in a single platform to provide simultaneous imaging and treatment functions.^1–3^ Agents that can provide such multiple functionalities via a simple, single-component design are particularly advantageous. This type of agent can be safer since fewer administrations are needed to achieve the desired outcomes. This so-called “one-for-all” methodology reduces overall fabrication complexities and improves the prospects for large-scale production and clinical use. For example, silver sulfide nanoparticles (Ag_2_S-NP) constitute a low-cost, highly biocompatible theranostic agent owing to their chemical inertness and remarkable optical properties, such as tunable near-infrared fluorescence (NIRF) emissions and photothermal conversion.^4^ However, previous Ag_2_S-NP formulations are synthesized in the hydrophobic phase using non-biocompatible precursors under harsh conditions (*e.g.*, thermal decomposition). They are mostly larger than the size threshold necessary for efficient renal clearance and are not optimized for both imaging and therapy.^5–7^ Most studies have only focused on their use in optical-based techniques,^8^ while multimodality imaging has become increasingly important for disease detection. Their photothermal effects can be limited due to the narrow band gap, weak NIR absorbance, and inefficient conversion of light to heat. Therefore, improvements to the synthesis and design of Ag_2_S-NP are needed to create a potentially translatable agent with enhanced theranostic performance.

Recently, we have found that sub-5 nm Ag_2_S-NP generate X-ray contrast well-suited for breast cancer detection since the K-edge of silver at 25.5 keV is ideal for the X-ray energy range used in mammographic imaging techniques, such as mammography, dual energy mammography, and tomosynthesis.^9, 10^ Moreover, these nanoparticles are efficiently eliminated through the renal system owing to their ultrasmall size.^11^ We have also developed size-tunable Ag_2_S-NP and other nanoparticle formulations in the aqueous phase using a microfluidic approach under ambient conditions, providing a facile and reproducible synthetic method.^12–14^ Herein, we present an innovative design approach to optimize the diagnostic and therapeutic functions of Ag_2_S-NP, especially for their optical properties. We have synthesized sub-5 nm Ag_2_S-NP with a narrow NIRF emission peak and encapsulated them in larger nanostructures made of a mixture of biodegradable poly[di(carboxylatophenoxy)phosphazene] (PCPP) and its poly(ethylene glycol) (PEG) containing graft copolymer (PEG-PCPP) to form AgPCPP nanoparticles using a microfluidic device. These polymers are effective drug delivery vehicles that can undergo hydrolytic degradation into physiologically benign hydrophilic products.^15–18^ This work represents the first to demonstrate the size tunability of these polymeric nanoparticles by modulating the amount of PEG-PCPP used in the formulation. We have found that the encapsulation of Ag_2_S-NP in PCPP greatly increases the NIR absorbance, resulting in enhanced photoconversion activities (both radiative and nonradiative emissions). We have shown that AgPCPP are biocompatible and biodegradable *in vivo* and are effective for multimodality detection with computed tomography (CT), NIRF (also image-guided surgery), and photoacoustic (PA) imaging. In addition, we have demonstrated that AgPCPP significantly inhibit breast tumor growth via photothermal therapy (PTT) and improve overall survival compared to free Ag_2_S-NP. Therefore, this “one-for-all” cancer theranostic agent can potentially be used for whole body imaging with CT, while enabling NIRF and PA image guided surgery and PTT localized to the tumor region, before gradually breaking down and releasing Ag_2_S-NP for excretion (Scheme 1),^11^ thus addressing concerns over long-term retention.

**Scheme 1.**
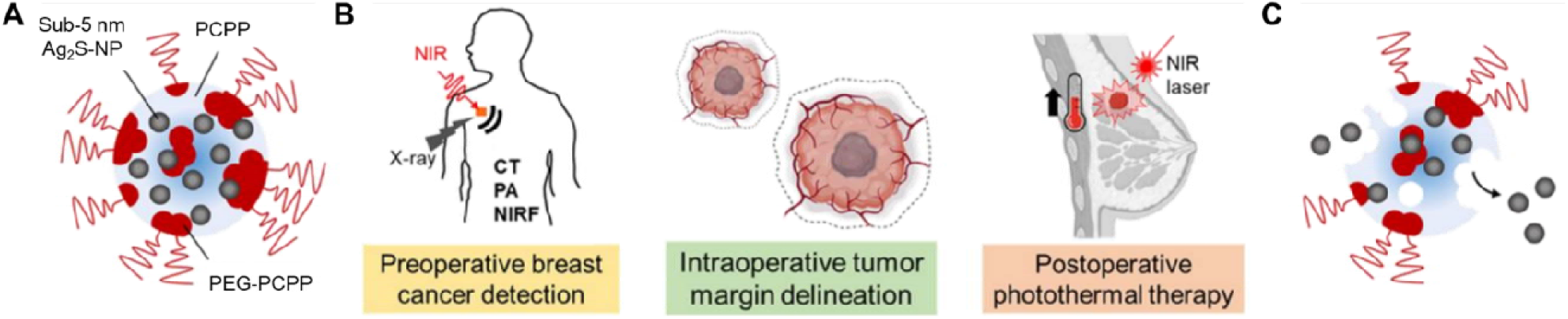
Schematic illustration of (a) AgPCPP nanoparticles, (b) their theranostic functions, and (c) their degradability and release of ultrasmall Ag_2_S-NP for excretion.

## Results

### Nanoparticle synthesis and characterization

Small core Ag_2_S-NP were synthesized in water by slowly adding sodium sulfide to a mixture containing 2-MPA and silver salt to trigger the nucleation and growth of silver cations into silver sulfide nanocrystals. The reaction was carried out at room temperature under ambient conditions. 2-MPA was used as a capping ligand to stabilize the silver sulfide cores and provide hydrophilicity and colloidal stability in biological media. Nanoparticles coated with this thiol ligand have previously demonstrated good biocompatibility with high renal clearance efficiency.^5, 19, 20^ The core diameter of Ag_2_S-NP was determined to be 1.7 ± 0.6 nm (Figure S1), while the hydrodynamic diameter was slightly larger (3.6 ± 1.3 nm). The zeta potential of Ag_2_S-NP was found to be -70 ± 20 mV owing to the negatively charged carboxylate groups of 2-MPA.

Ag_2_S-NP were encapsulated into spermine cross-linked ionotropic hydrogel nanoparticles of PCPP and PEG-PCPP using a microfluidic device. The resulting AgPCPP nanoparticles are tunable in size. The size of AgPCPP (0.5 mg Ag per mg total PCPP) could be controlled by varying the amount of PEG-PCPP copolymer used in the synthesis, as evidenced by TEM and SEM (Figure 1A), from 40 to 300 nm. The average diameter of AgPCPP decreased with an increasing content of PEG-PCPP in a nonlinear fashion (Figure 1B). Our finding is consistent with previous reports, whereby the inclusion of a block copolymer limits and controls the growth of PCPP nanoparticles.^15, 21^ For subsequent experiments, we used AgPCPP nanoparticles that were synthesized with 10% PEG-PCPP in the formulation (82.1 ± 12.8 nm in diameter), *i.e.,* 0.1 mg PEG-PCPP, and 0.5 mg Ag per mg of total PCPP polymers. This formulation was chosen since its size was similar to current FDA approved nanodrugs, such as Doxil (90 nm PEGylated liposomal doxorubicin), that are long circulating with high tumor accumulation.^22^

**Figure 1.**
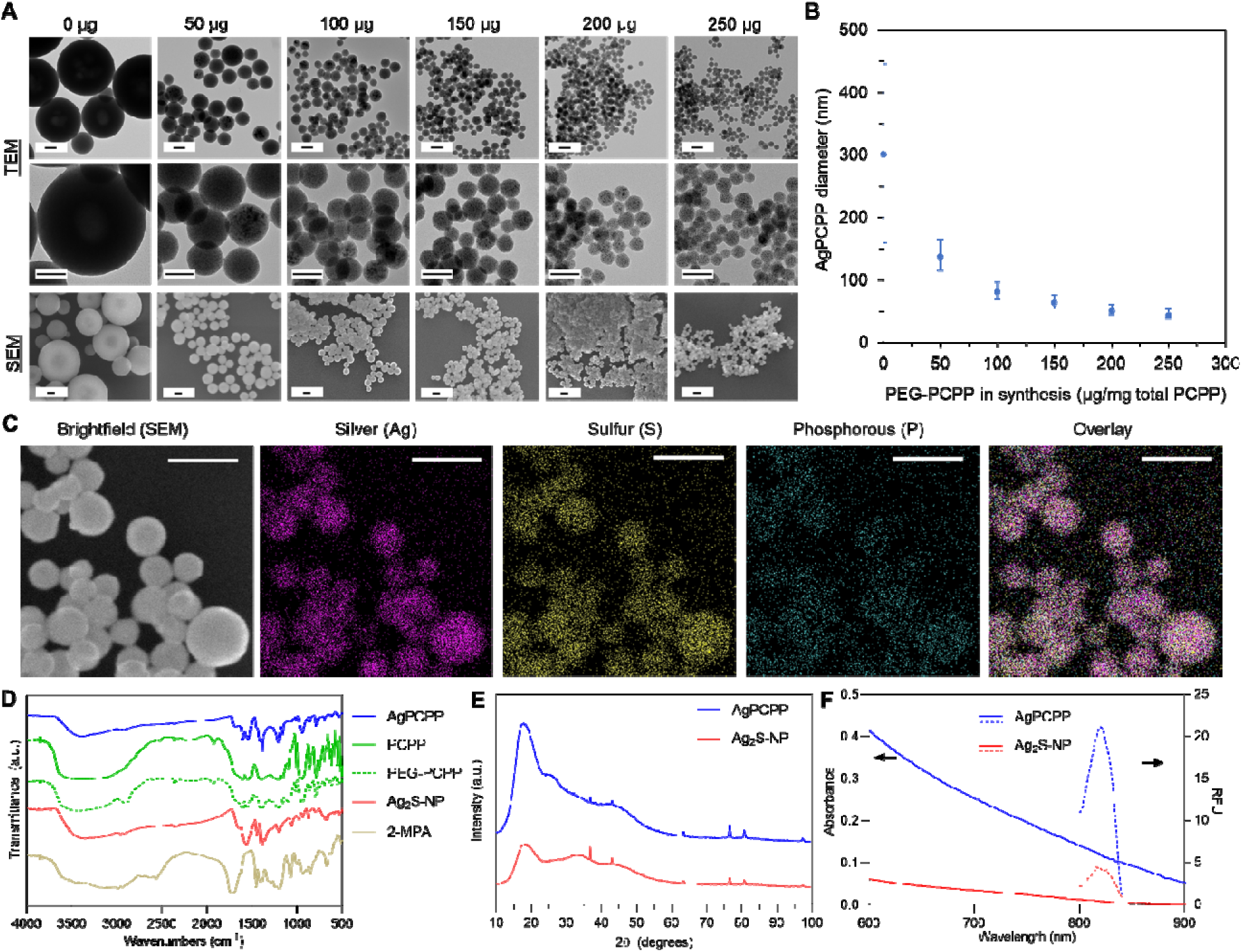
Characterization of AgPCPP nanoparticles. (A) AgPCPP formulated with varying amounts of PEG-PCPP (presented as µg PEG-PCPP per mg of total polymers). Micrographs were acquired by transmission electron microscopy (TEM – top two rows) and scanning electron microscopy (SEM – bottom row). Scale bar = 100 nm. (B) Effect of PEG-PCPP on AgPCPP nanoparticle size. Error bars are one SD. (C) Elemental mappings of AgPCPP nanoparticles via energy dispersive X-ray spectroscopy (EDX). Scale bar = 250 nm. (D) Infrared spectra of AgPCPP nanoparticles and their individual components via Fourier transform infrared spectroscopy (FT-IR). (E) Powder X-ray diffraction (XRD) patterns and (F) UV-visible absorption (solid lines) and near-infrared fluorescence (dashed lines, ex: 700 nm) spectra of AgPCPP and Ag_2_S-NP.

Analysis of a single AgPCPP nanoparticle via energy dispersive X-ray spectroscopy (EDX) revealed several elemental peaks, including phosphorous, silver and sulfur, with an expected 2:1 Ag:S stoichiometry (Figure S2). The phosphorous signal arises from the polyphosphazene polymer backbone. EDX mappings further confirmed the successful incorporation of Ag_2_S-NP into the polymer matrix, as all three major elements were detected in every AgPCPP nanoparticle (Figure 1C). Fourier transform infrared spectroscopy also confirmed the inclusion of both PCPP polymer types in AgPCPP nanoparticles owing to the characteristic peaks at 1550-1700 cm^-1^ (C = O stretching). The peak at 2500 cm^-1^ is absent, which corresponds to the formation of Ag-thiol bond and hence the presence of Ag_2_S-NP in AgPCPP nanoparticles (Figure 1D). Moreover, the powder XRD patterns of AgPCPP nanoparticles are broad with few distinctive peaks due to the ultrasmall size of Ag_2_S-NP but match well with the monoclinic crystalline structure of Ag_2_S (Figure 1E). AgPCPP nanoparticles exhibit a broad UV-visible absorption band (Figure 1F), which is typical of Ag_2_S-NP.^23^ The broadband absorbances in the NIR region suggest that these nanoparticles could achieve photothermal conversion under NIR laser irradiation.^24^ Interestingly, we observed an aggregation induced red shift in the absorbance of AgPCPP nanoparticles. Furthermore, under excitation at 700 nm, we found that both Ag_2_S-NP and AgPCPP nanoparticles have a narrow fluorescence emission peak at 820 nm (Figure 1F). Notably, AgPCPP nanoparticles have much stronger NIR fluorescence intensity than free Ag_2_S-NP at the same silver concentration. Therefore, encapsulation into larger polymeric nanoparticles can potentially enhance the optical properties of the small metal core nanoparticles.

### *In vitro* biocompatibility and PTT performance

Ag_2_S-NP and AgPCPP nanoparticles were incubated with endothelial cells, hepatocytes, kidney epithelial cells, and breast cancer cells for 24 hours to assess their cytotoxicity via MTS assay. Incubation with Ag_2_S-NP and AgPCPP did not affect cell viability (>99%) at concentrations up to 1 mg Ag/ml (Figure S3). Both nanoparticle types were found to be biocompatible at each concentration with no statistically significant differences compared to control. The results attest to the non-toxic properties of PCPP polymers^25^ and the inert and insoluble nature of silver sulfide.^9, 11^

To evaluate the photothermal conversion activities, we measured the temperature elevation by Ag_2_S-NP and AgPCPP nanoparticles when irradiated under an 808 nm laser. AgPCPP increased the temperature of the solution by 40 °C during 10 minutes of irradiation at a power density of 2 W per cm^2^ and a silver concentration of 0.5 mg per ml (Figure 2A). However, Ag_2_S-NP raised the solution temperature by only 10 °C under the same conditions. The photothermal conversion efficiency of AgPCPP was calculated to be 25.2%,^26^ which is comparable to other reported structures, such as gold nanorods and ICG-loaded micelles.^27–30^ We then evaluated the photostability of AgPCPP under four irradiation/cooling cycles (Figure 2B). We found that AgPCPP nanoparticle solution reproducibly reached 99.9% of the temperature (60 °C) achieved the first time, indicating excellent stability of these nanostructures. Their structure and size remained unchanged (79.5 ± 11.5 nm) after repeated irradiation cycles, as evidenced by TEM (Figure S4), suggesting that the polymers were not affected or degraded by photothermal heating. Additionally, we found that temperature elevation by AgPCPP is dependent on the concentration of silver payloads (Figure S5A) and laser power density (Figure S5B). These results suggest that incorporation of Ag_2_S-NP into larger polymeric nanoparticles effectively strengthens their photothermal properties.

**Figure 2.**
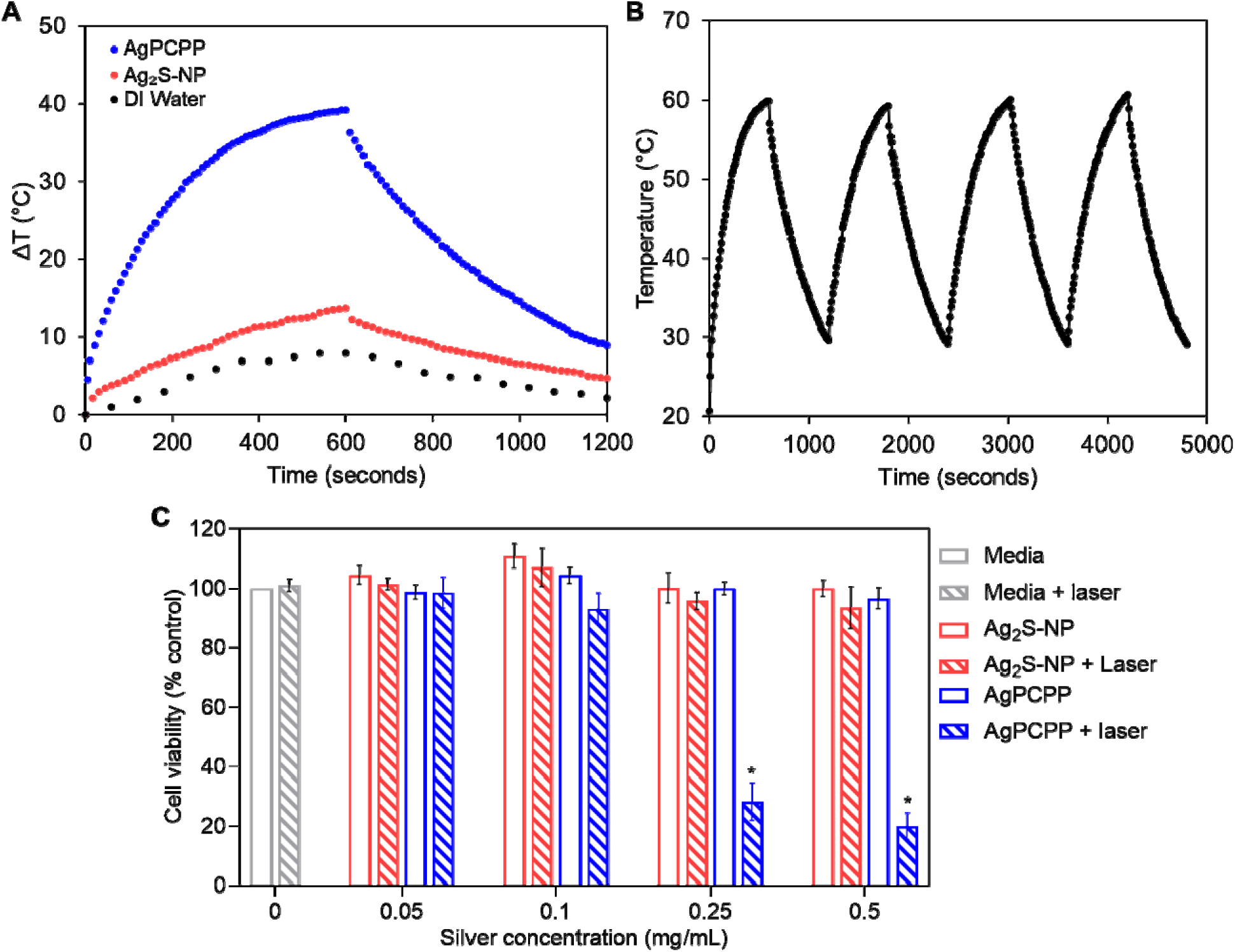
Photothermal activity and killing effect of Ag_2_S-NP and AgPCPP. (A) Heating and cooling curves of AgPCPP nanoparticles, free Ag_2_S-NP, and DI water under 10 minutes of irradiation (808 nm, 2 W/cm^2^). (B) Photostability of AgPCPP under four cycles of irradiation/cooling for a total duration of 20 minutes (808 nm, 2 W/cm^2^). (C) Viability of MDA-MB-231 breast cancer cells incubated with varying concentrations of Ag_2_S-NP and AgPCPP in darkness and under laser irradiation (808 nm, 1.5 W, 5 minutes). \**p* < 0.05 compared to media only control group (one-way ANOVA with Tukey’s multiple comparisons test).

We then examined the photothermal heating effects of Ag_2_S-NP and AgPCPP nanoparticles on the viability of MDA-MB-231 breast cancer cells. Cells were incubated with increasing concentrations of Ag_2_S-NP or AgPCPP, after which the cells were subjected to 5 minutes of laser irradiation at a power of 1.5 W. As shown in Figure 2C (hatched color bars), treatment with media or free Ag_2_S-NP in the presence of laser irradiation did not adversely affect cell viability. A slight reduction in viability (94 ± 7%) was observed from free Ag_2_S-NP at the highest silver concentration. This is not statistically significant when compared to non-irradiated control. However, laser irradiation treatment with AgPCPP at 0.25 mg/ml and above resulted in significant decreases in cell viability. This finding is consistent with the fact that AgPCPP can elevate temperature more effectively than free Ag_2_S-NP, thus resulting in greater photothermal killing. Therefore, AgPCPP can produce potent photothermal therapeutic effects and have potential for eliminating laser irradiated tumor tissues.

### Multimodality phantom imaging

We first assessed the potential of AgPCPP nanoparticles as a photoacoustic based imaging agent. The PA images were acquired using a laser frequency of 700 nm and a PA gain of 24 dB (Figure 3A). PA signals were generated based on thermoelastic expansion from Ag_2_S metal cores. AgPCPP generated a significantly higher PA signal than free Ag_2_S-NP due to the aggregation induced red shift in their UV-visible absorption spectrum (Figure 3B). AgPCPP produced contrast that is linearly correlated with concentration (R^2^ = 0.99). However, Ag_2_S-NP samples did not produce a significant PA signal, likely due to their low absorption of NIR light. Next, we investigated the NIRF optical contrast properties using an IVIS Spectrum imaging system with a 710/820 nm filter pair (Figure 3C). For both Ag_2_S-NP and AgPCPP formulations, the fluorescence intensity observed becomes greater with increasing silver concentrations. The fluorescence intensity begins to plateau at a concentration of 2 mg Ag/ml (Figure 3D). Moreover, AgPCPP produced higher NIR fluorescence signal than Ag_2_S-NP over the same range of silver concentrations. Interestingly, the aggregation of Ag_2_S-NP into the polymer matrix did not induce quenching of fluorescence signal from Ag_2_S-NP. Lastly, we examined the suitability as a fluorescence image-guided surgery agent using the Vet-FLARE system. Figure 3E shows the resulting brightfield, NIR 700, NIR 800 and merged (with pseudo colors) images. As expected, no contrast was observed in the NIR 700 images since both Ag_2_S-NP and AgPCPP have an emission peak above 800 nm. In the NIR 800 images, AgPCPP showed significantly higher (by three-fold) fluorescence signal intensity than free Ag_2_S-NP over the same concentration range (Figure 3F). The intensity starts to plateau at the concentration of 2 mg Ag/ml, which is similar to the finding from IVIS Spectrum imaging experiments.

**Figure 3.**
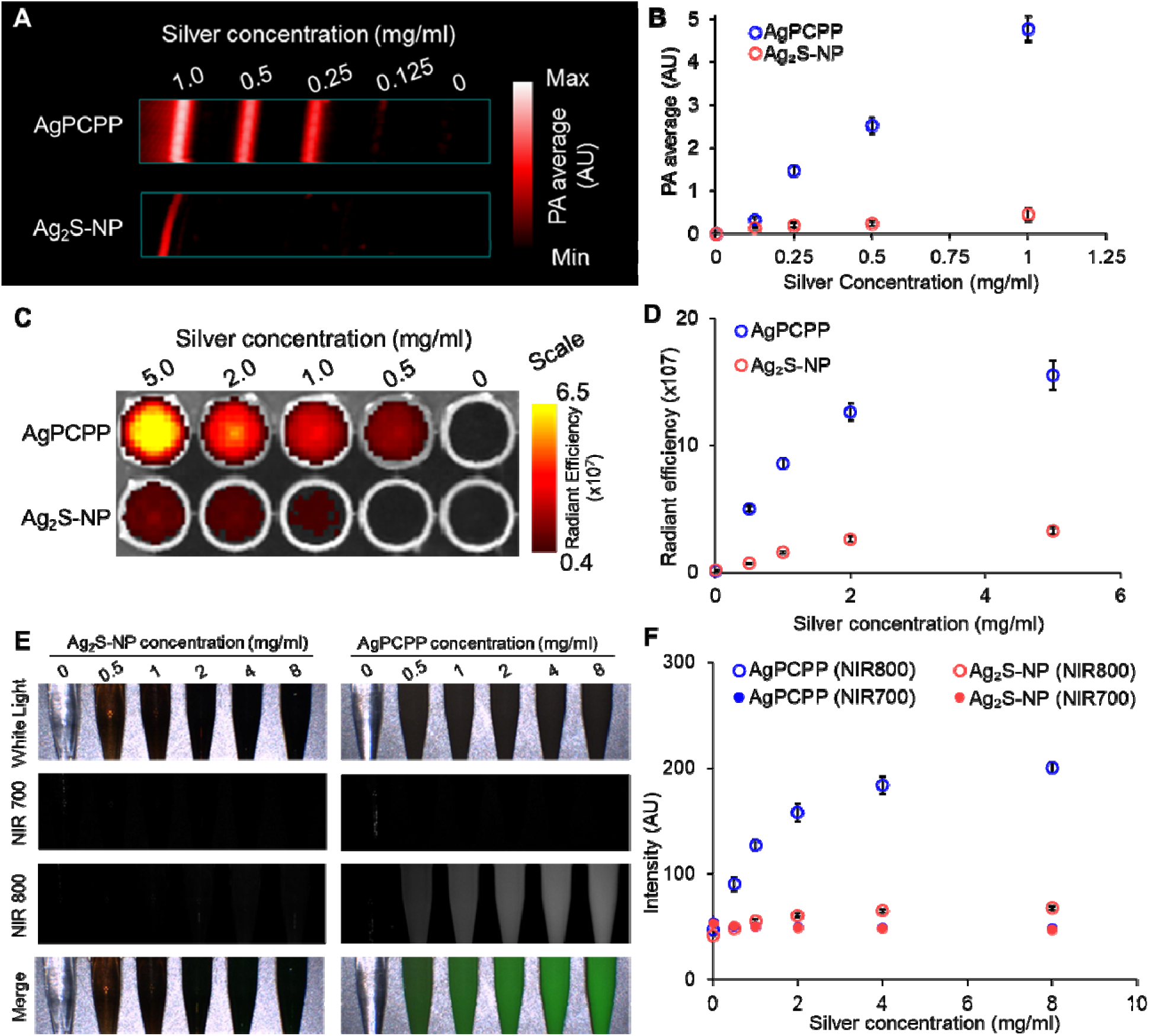
Optical-based contrast properties of Ag_2_S-NP and AgPCPP. (A) Phantom images from a PA imaging system. (B) Relationship between PA contrast and Ag concentration. (C) NIRF phantom image of both nanoparticle formulations. Scale bar in units of radiant efficiency (x10^7^ (p/s/cm^2^/sr)/μW/cm^2^). (D) Relationship between NIRF intensity and Ag concentration. (E) Color, NIR 700, NIR 800 and merged images from an intraoperative, imaged-guided surgery system. (F) Quantification of fluorescence signal intensity from both NIR imaging channels and both nanoparticle types at a range of silver concentrations. Some error bars are smaller than the data point markers, while some markers overlap with and mask other markers, and are thus not visible.

We also determined the CT contrast properties of Ag_2_S-NP and AgPCPP using both a µCT scanner and a clinical CT system. A µCT scanner operated at a tube voltage of 50 kVp was used to scan phantom samples of both formulations (Figure 4A) and control materials, including silver salt and iodine. All materials produced CT contrast that is linearly correlated between attenuation and concentration as expected, with an R^2^ value > 0.99 in each case (Figure S6). We found that silver nitrate, Ag_2_S-NP and AgPCPP have statistically significantly higher attenuation rates than iopamidol (Figure 4B). The attenuation rates among all silver-based contrast materials were not found to be statistically significantly different. Furthermore, the same phantom samples were scanned in a clinically relevant anthropomorphic phantom body that closely represents a patient’s thorax using a clinical CT scanner at four different tube voltages. Axial plane images of the phantom body containing both nanoparticle formulations at tube voltage of 80 kVp are shown in Figure 4C. An increase in CT contrast was observed with increasing concentrations of the payload. As expected, the attenuation rates of all silver-based contrast materials are not significantly different among each other but are lower and statistically significantly different than that of iopamidol at each tube voltage (Figure 4D). For all agents, the highest CT attenuation rate occurred at 80 kVp and the rates decreased as the tube voltage increased to 140 kVp. On the contrary, silver has a higher attenuation rate than iodine in µCT imaging, which can be attributed to the lower kVp used in µCT, thus bringing the mean photon energy closer to silver’s K-edge. Notably, sufficient contrast was acquired with AgPCPP at a minimum concentration of 1 mg Ag/ml in all modalities (*i.e.*, optical and X-ray imaging). AgPCPP nanoparticles are therefore well suited for multimodality imaging applications owing to balanced contrast production. Overall, AgPCPP have potential use as an optical agent with enhanced performance, while providing good X-ray CT contrast .

**Figure 4.**
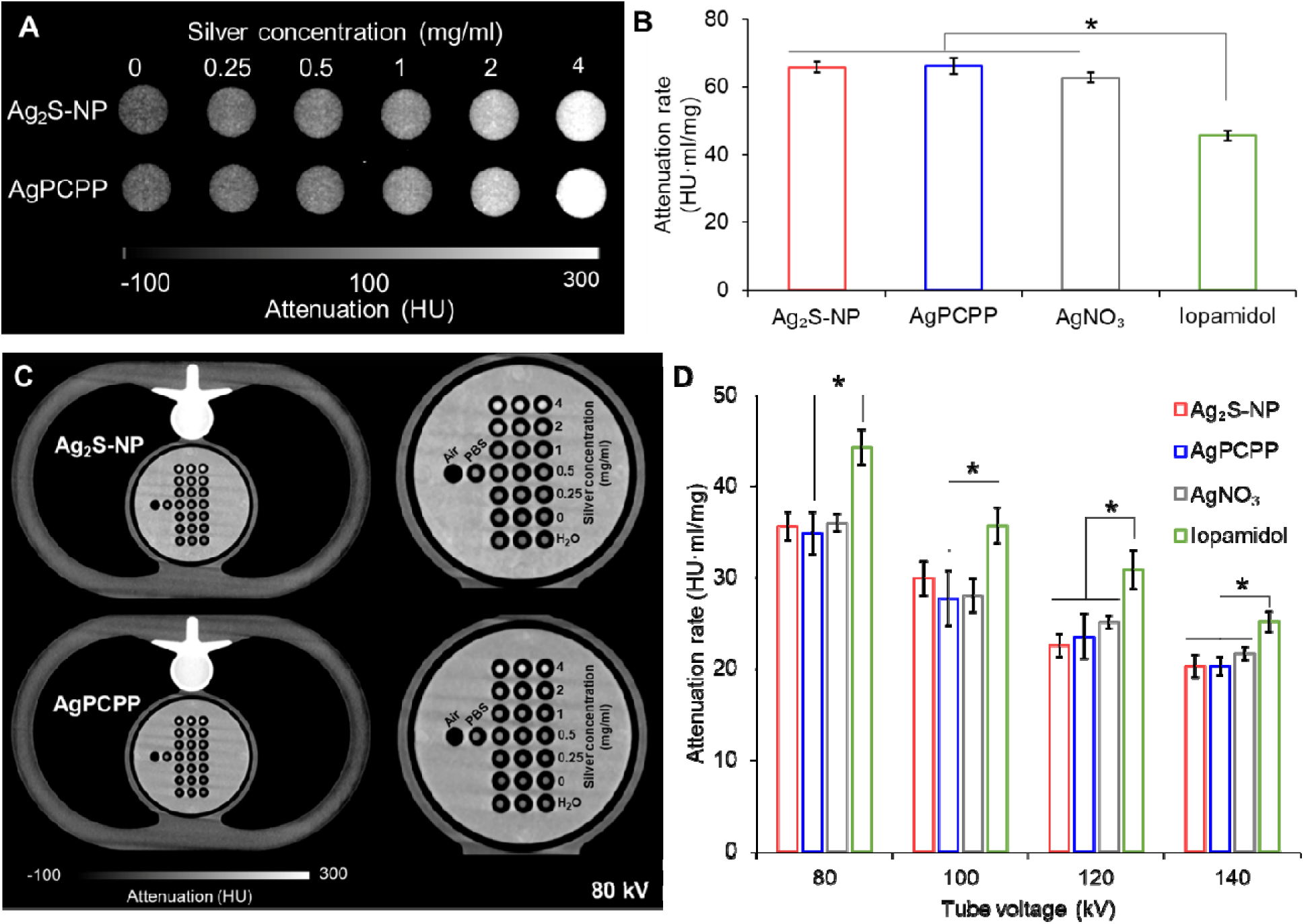
*In vitro* CT contrast generation of Ag_2_S-NP and AgPCPP. (A) Phantom images from a MILabs µCT scanner. (B) Attenuation rates of both nanoparticle formulations and control materials. (C) Phantom images taken with a clinical CT scanner. (D) Attenuation rates of both nanoparticle formulations and control materials at different tube voltage settings. Data are presented as mean ± SD. Error bars are one SD. \**p* < 0.05 (one-way ANOVA with Tukey’s multiple comparisons test).

### *In vitro* release of Ag_2_S-NP

We evaluated the release of Ag_2_S-NP from their polymeric components by incubating AgPCPP nanoparticles in 10% serum in PBS at 37 °C. We found that approximately 88% of Ag_2_S-NP were released from AgPCPP nanoparticles in 7 days (Figure 5A), indicating the biodegradability of these nanoparticles and effective release of the payloads. TEM (Figure 5B) and SEM (Figure 5C) images of the pellet collected at the end of the incubation period showed the breakdown and morphological deformation of AgPCPP nanoparticles, as well as the release of small core Ag_2_S-NP.

**Figure 5.**
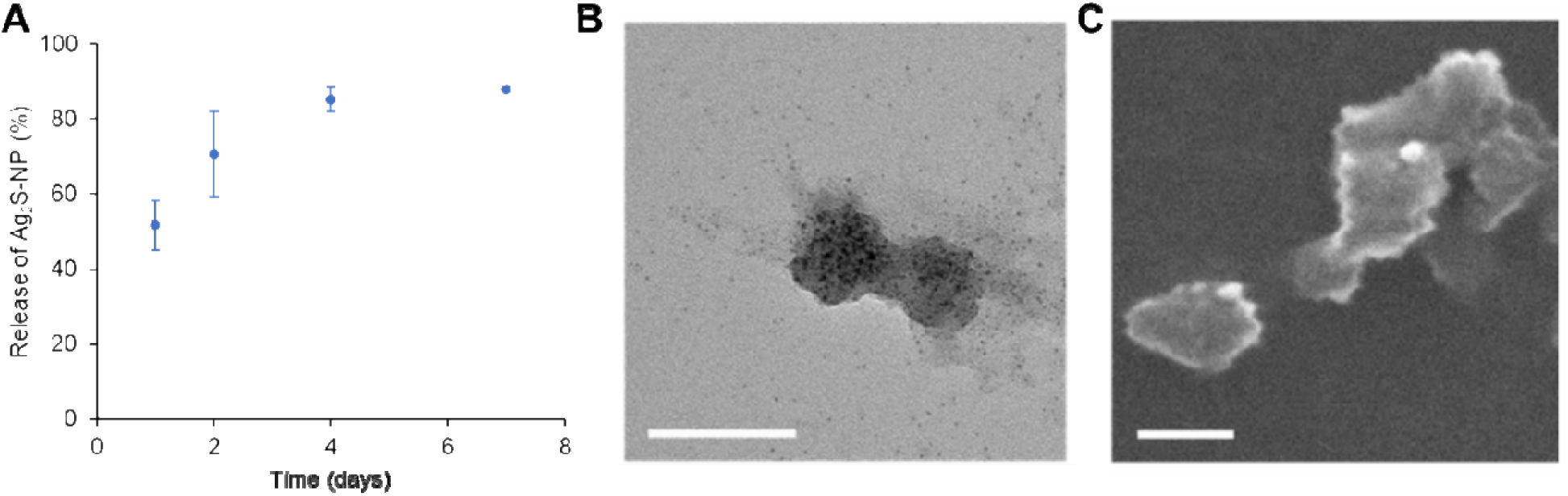
*In vitro* biodegradability of AgPCPP nanoparticles. (A) Payload release from AgPCPP nanoparticles. Data are presented as mean ± SD. Error bars are one SD. (B) TEM and (C) SEM images of partially degraded AgPCPP nanoparticles after 7 days of incubation. Scale bar = 100 nm.

### *In vivo* multimodality imaging

A murine model of breast cancer was used to investigate the applicability of AgPCPP nanoparticles as an effective contrast agent for *in vivo* multimodality tumor detection. As shown in Figures 6A and S7, the representative images showed punctate CT contrast enhancement at the injection site (in red dashed circles). Significantly higher CT attenuation was found in the tumors at all post-injection time points compared to the pre-injection scan (Figure 6B). No significant difference in CT attenuation was found among the post-injection time points. Similar results were obtained from *in vivo* NIRF imaging studies as presented in Figure 6C. The average NIRF signal intensity in the tumors significantly increased after injections of AgPCPP nanoparticles (Figure 6D). However, the signal decreased over the one-week time period, and the differences were found to be statistically significant. The signal loss could be attributed to the tumor growth that impacted light penetration, as well as the degradation of polymeric cores due to hydrolysis. Moreover, AgPCPP nanoparticles improved the visualization of tumors by PA imaging as shown in Figure 6E. The signal from AgPCPP nanoparticles was spectrally distinguishable from background PA signal provided by surrounding tissue, and their PA spectra from both post-injection time points followed a similar pattern as that obtained from *in vitro* PA phantom imaging (Figure 6F). Therefore, our data demonstrate that AgPCPP nanoparticles can readily enhance tumor contrast and serve as a potent imaging agent for a variety of modalities.

**Figure 6.**
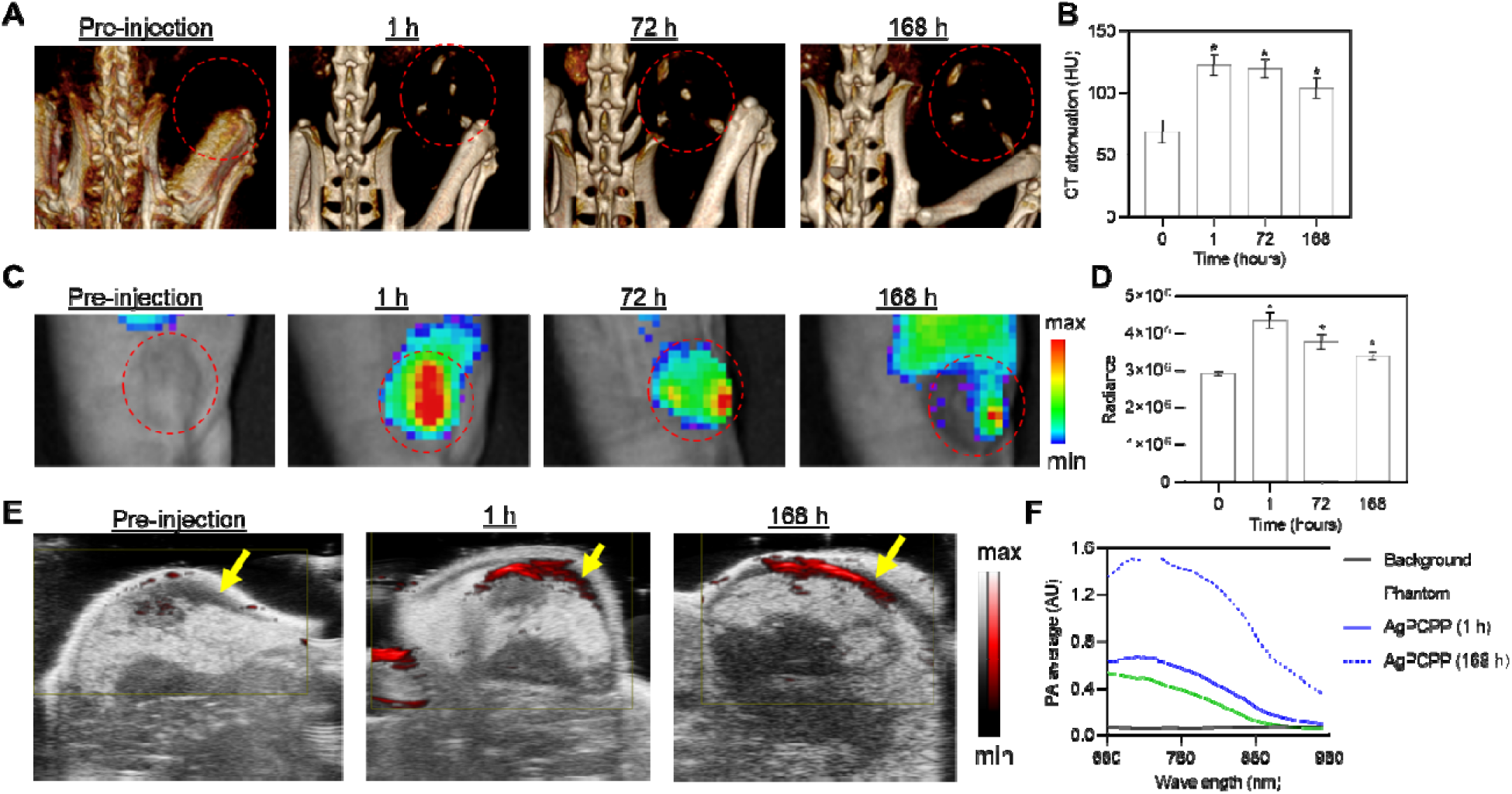
*In vivo* multimodality breast tumor imaging with AgPCPP nanoparticles. (A) Representative 3D volume rendered CT image reconstructions. The images are shown with a window level of 475 HU and window width of 650 HU. Red dashed circles indicate the tumor site. (B) Quantification of the CT attenuation in the tumors. (C) Representative NIRF images. Red dashed circles indicate the tumor site. (D) Quantification of the NIRF signal intensity in the tumors (radiance = photons/second/cm^2^/sr). (E) Representative ultrasound images with PA signal overlay. Yellow arrows indicate the tumor site. (F) PA spectra of background and injection site within the NIR wavelength range. \**p* < 0.05 compared to pre-injection scan (one-way ANOVA with Tukey’s multiple comparisons test).

### *In vivo* photothermal heating and therapy

A PTT model was used to evaluate the antitumor efficacy of Ag_2_S-NP and AgPCPP nanoparticles. Mice with established breast tumors were treated with either saline or nanoparticles, and a subset of mice received laser irradiation. As shown in Figure 7A, an increase in tumor temperature was observed for all groups during irradiation. Note that the infrared thermographic camera detects only the surface temperature and is not capable of characterizing the thermal environment within tissues. Importantly, AgPCPP nanoparticles achieved the highest peak tumor temperature (51 °C) compared to those achieved by the control (38 °C) and Ag_2_S-NP (44°C) groups (Figure S8). After 30 minutes of irradiation, AgPCPP nanoparticles raised the tumor temperature relative to overall body temperature by 17°C, while saline and Ag_2_S-NP raised the tumor temperature by 6 °C and 11 °C, respectively (Figure 7B). The steady temperature elevation during laser irradiation also indicates the structural stability and photostability of AgPCPP nanoparticles. These results are consistent with the outcome from *in vitro* photothermal ablation studies. Moreover, their potent photothermal conversion activity should lead to ideal therapeutic efficacy. Tumor growth of control and treatment groups were tracked for 14 days after PTT (Figures 7C and S9). Tumor growth was greatly reduced in mice that received the therapeutic combination of AgPCPP nanoparticles with laser irradiation compared with all other groups. The average tumor size regressed below pre-treatment volume within 4 days of therapy and remained similar to initial volume for a total of 10 days after treatment. After that, tumor regrowth was observed, likely due to incomplete eradication of deep-seated tumor tissues as a result of limited light penetration. No other group displayed tumor regression or reached pre-therapy volume. On Day 14, tumors treated with AgPCPP nanoparticles and laser irradiation were markedly much smaller in overall size compared to the other groups (Figure S10). Furthermore, we found a significant improvement in survival for mice receiving AgPCPP nanoparticles and laser irradiation, as none of them met the criterion for sacrifice, while all other groups had reduced survival by the end of the study (Figure 7D). Additionally, the body weights of all groups were not significantly different between pre- and post-treatment (Figure S11). The stability in weights suggests that the mice were in good health and were not impacted by the treatments despite continued tumor growth in most groups. Therefore, AgPCPP nanoparticles along with laser irradiation exhibit potent tumor hyperthermia, good photostability and biocompatibility, and remarkable antitumor effects.

**Figure 7.**
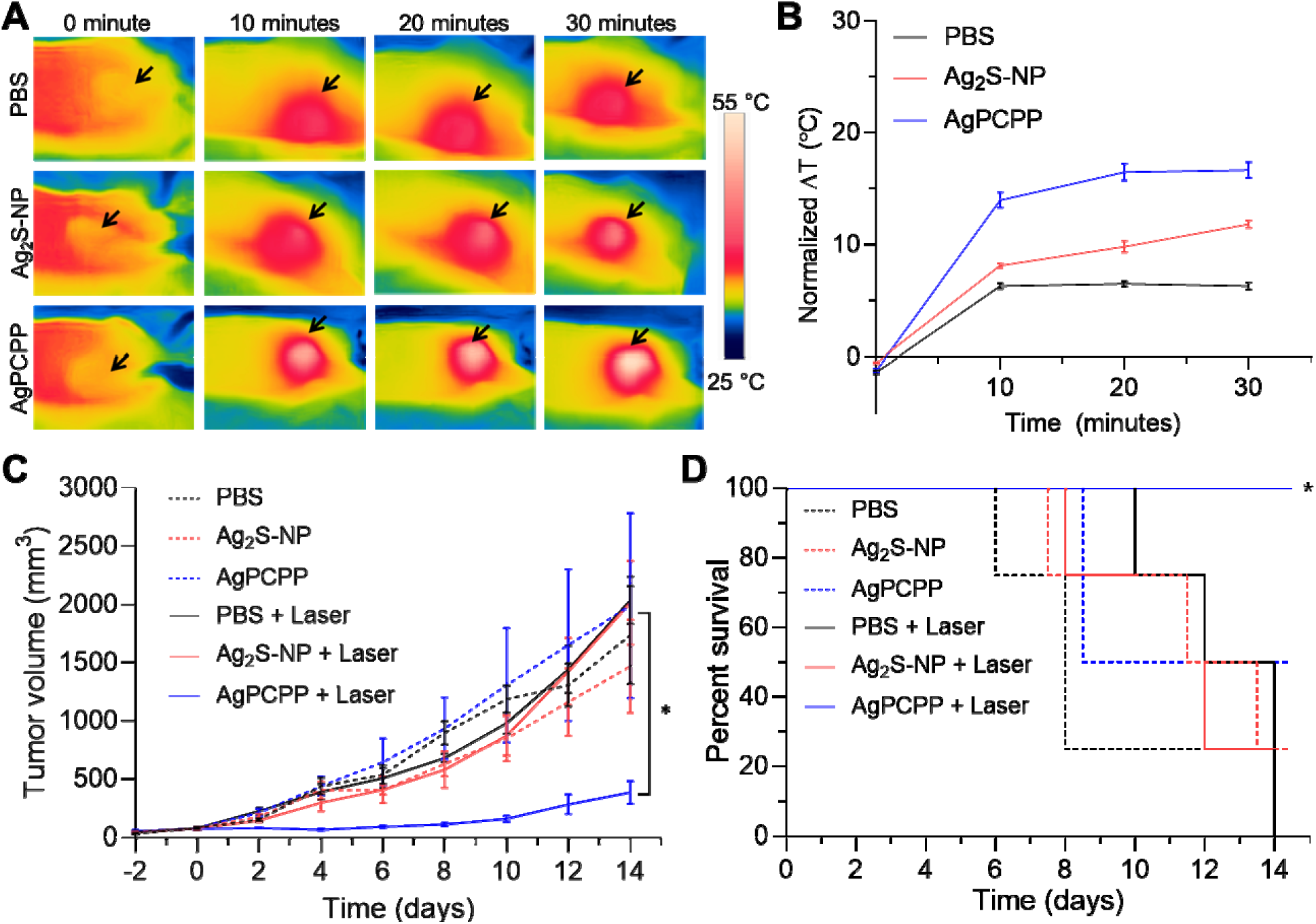
*In vivo* photothermal ablation and treatment. (A) Infrared thermography and (B) temperature elevation of tumors treated with saline, Ag_2_S-NP, and AgPCPP nanoparticles during 30 minutes of laser irradiation. Black arrow indicates the tumor site. (C) Tumor growth of mice treated with saline, Ag_2_S-NP, and AgPCPP nanoparticles with or without 30 minutes of laser irradiation. day 0 = tumor injection and laser treatment. \**p* < 0.05 (two-way ANOVA with Bonferroni’s multiple comparisons test). (D) Kaplan-Meier curves demonstrating animal survival for 14 days after photothermal treatment. \**p* < 0.05 (log-rank test).

### *In vivo* toxicity

We assessed *in vivo* safety of AgPCPP nanoparticles via histological examination of major organs at 1 day and 7 days post injection. As shown in Figure 8A, micrographs of H&E stained tissues did not show any evident microscopic structural changes or abnormal pathologies in comparison with the control population. As presented in Figure 8B, the differences in the serum biomarker levels between the control and treatment groups were not statistically significant across all biomarkers analyzed. Hence, the liver and kidney functions were not impacted by the retention of these nanoparticles. Together, the histopathology and serum biochemistry panels indicated that no signs of acute toxicity or adverse effects were observed in treated mice up to a week of exposure. Therefore, these preliminary results supported the *in vivo* safety of AgPCPP nanoparticles.

**Figure 8.**
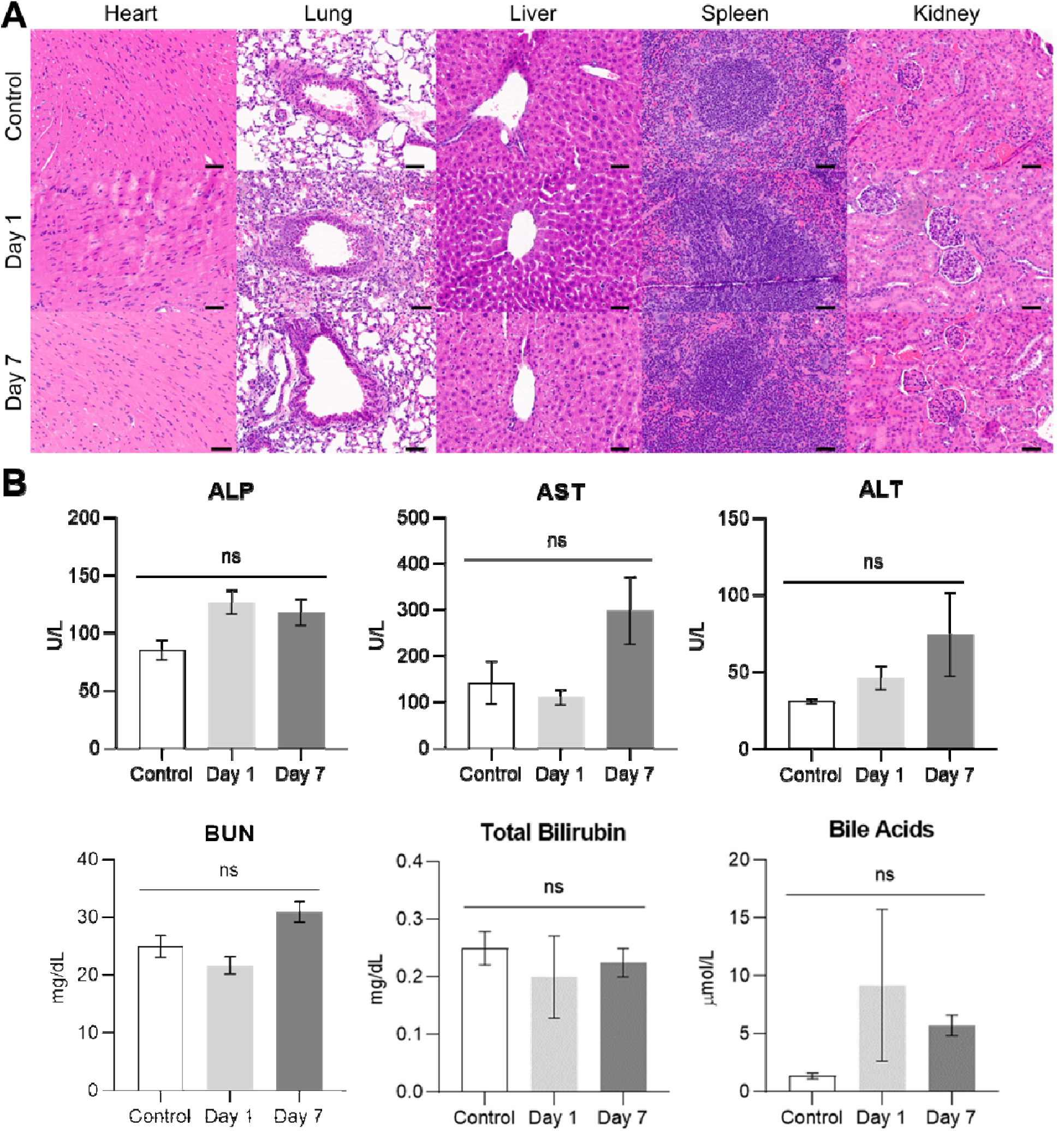
*In vivo* toxicology. (A) Representative H&E stained micrographs of major mouse organs after injection with either saline or AgPCPP nanoparticles at different time points post-injection. Scale bar = 50 µm. (B) Serum biomarker levels of liver and kidney functions from mice injected with either saline or AgPCPP nanoparticles at different time points post-injection. ALP = alkaline phosphatase, AST = aspartate aminotransferase, ALT = alanine transaminase, BUN = blood urea nitrogen. ns = not significant (one-way ANOVA with Tukey’s multiple comparisons test).

### Biodistribution and *in vivo* biodegradation

We first evaluated the tumor uptake between Ag_2_S-NP and AgPCPP nanoparticles. At 24 hours after intravenous injection, tumor-bearing mice were euthanized, and their organs and tumors tissues were collected for biodistribution analysis via ICP-OES. The results indicated that higher amounts of AgPCPP nanoparticles were found in the liver and spleen compared to Ag_2_S-NP (Figure S12A), while more Ag_2_S-NP were detected in the kidneys, indicating that these small core nanoparticles were being filtered through the renal system. The degree of tumor accumulation was higher for AgPCPP nanoparticles compared to Ag_2_S-NP (Figure S12B), suggesting that the combination of surface PEGylation and large particle size prolonged blood circulation and facilitated uptake via the enhanced permeability and retention effect. We further investigated the long-term biodistribution of AgPCPP nanoparticles in non-tumor-bearing mice. The results showed a statistically significant reduction in nanoparticle accumulation in the liver and spleen at 7 days post-injection as opposed to 1-day post-injection (Figure 9A). This finding demonstrated the biodegradability of AgPCPP nanoparticles *in vivo* and the release and clearance of small core nanoparticles over time. Moreover, we examined the liver tissues *ex vivo* via TEM and found that AgPCPP nanoparticles were mainly localized in the Kupffer cells or macrophages in the liver. Their morphology and structure were mostly intact at 1-day post-injection but were mostly deformed at 7 days post-injection (Figure 9B). This observation further supported the long-term biodistribution results and confirmed the dissociation of the polymeric cores *in vivo*.

**Figure 9.**
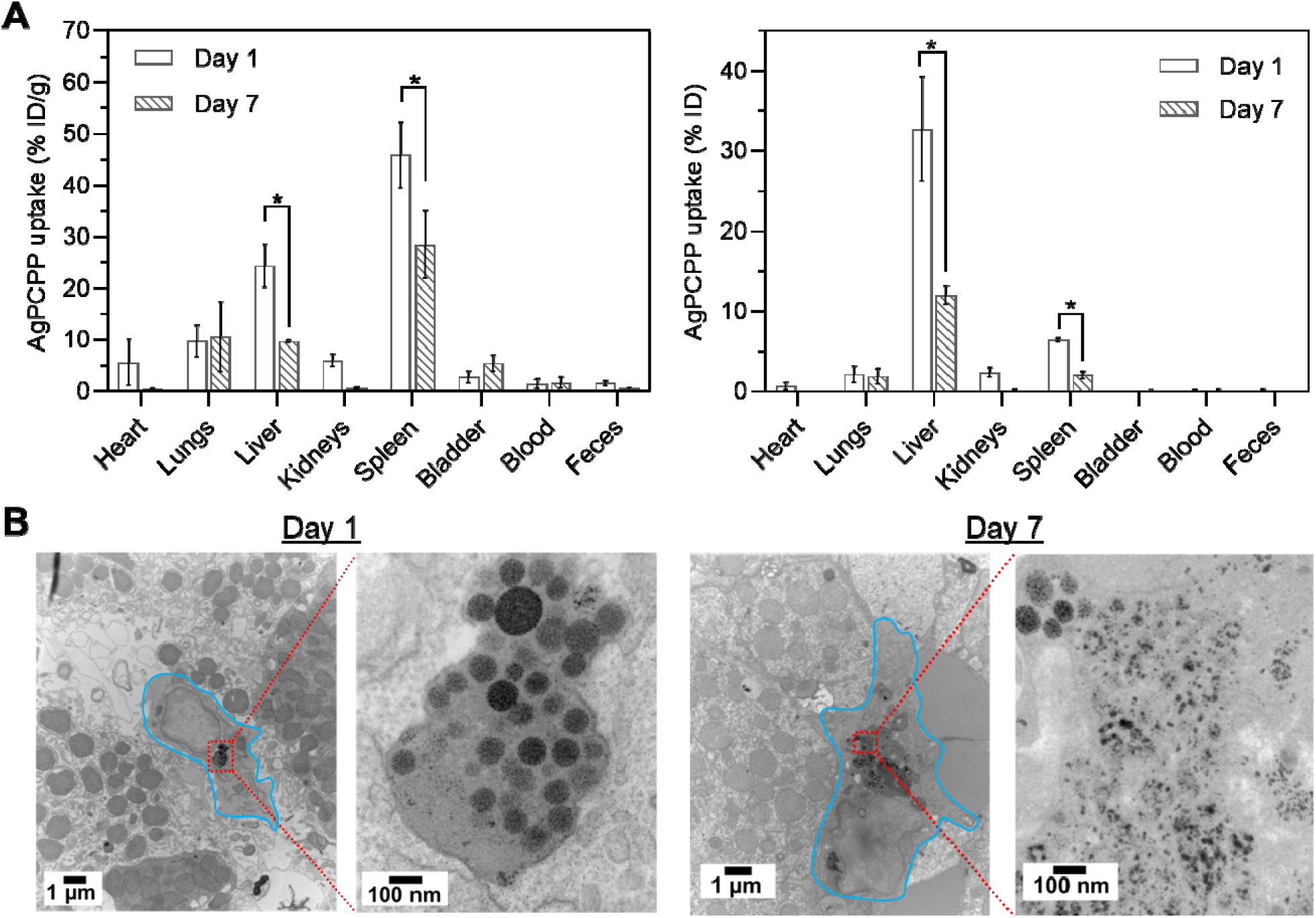
Biodistribution and biodegradation of AgPCPP nanoparticles. (A) Biodistribution in units of % ID/g and % ID at different time points post-injection. ID = injected dose. \**p* < 0.05 (two-way ANOVA with Bonferroni’s multiple comparisons test). (B) Representative TEM micrographs of liver tissues collected at different time points post-injection. Kupffer cells are outlined in blue, while nanoparticles are indicated by a red dashed box.

## Discussion

In this study, we have demonstrated effective synthetic control over nanoparticle size, which is one of the important factors in designing nano-based agents tailored for specific biomedical applications. We have observed significant improvements to the optical and photoconversion characteristics of small Ag_2_S-NP after being incorporated into large polymeric nanoparticles. AgPCPP, or aggregates of Ag_2_S-NP, exhibit desirable tumor contrast enhancement with various modalities, including CT, NIRF, and PA imaging. Notably, the balanced contrast production of AgPCPP across multiple different modalities allows for simplified administration and reduced dosing. AgPCPP also show stronger PTT performance as well as greater anti-tumor effects compared to free Ag_2_S-NP. Overall, this is an innovative approach to improving an agent’s diagnostic and therapeutic efficacy. It is worth nothing that our nanoplatform can be used for imaging and treating other types of cancer, in addition to breast cancer as demonstrated in this work. AgPCPP rely on passive targeting and exhibit desired X-ray and optical contrast properties, as well as photothermal effects. These properties may also be applicable to other types of cancer.^31^ PTT cancer treatments are limited by the depth penetration of light in tissue, which is usually estimated at about 1 cm. However, for many cancers, including breast cancer, surgery is frequently part of the treatment process, which provides an opportunity to directly irradiate the tumor cavity. On the other hand, optical fibers can be advanced into orifices such as the trachea or rectum or inserted into tumor directly through the skin, thereby allowing irradiation of deeply seated tumors.^32^

Previous reports have utilized a cationic PEG-poly(L-lysine) block copolymer (PEG-PLL) to adjust the PCPP nanoparticle size.^12, 13^ Our platform herein is advantageous over previous formulations since it avoids the use of potentially reactogenic cationic non-PCPP polymers, which could further streamline the production and regulatory approval processes.^33^ Nanoparticle size modulation by PEG-PCPP is likely due to the stabilizing steric hindrance effect of PEG chains, which reduce interpolymer aggregation and lead to the formation of particles containing fewer polymer chains.^15^ The degradation profiles of AgPCPP nanoparticles can also be controlled via selecting polymers with different molecular weights or modifying the content of PEG groups. The assembly of AgPCPP and stability of the resulting nanoparticles are mainly determined by electrostatic interactions of negatively charged PCPP, PEG-PCPP, and Ag_2_S-NP with positively charged cross-linker – spermine. Therefore, it can be envisioned that varying the crosslinking density or using hydrolytically degradable cross-linkers with higher content of amino groups, such as previously described hydrolytically degradable PEGylated polyphosphazenes containing amino groups,^15^ can further increase *in vivo* half-life of nanoparticles.

Other groups have developed imaging agents based on aggregates of fluorescent quantum dots. For example, Fan *et al*. encapsulated hydrophobic cadmium-based quantum dots in a polymer matrix material, forming nanoparticles with a mean diameter of 27 nm.^34^ Chen *et al*. used siloxane surfactant to encapsulate quantum dots of the same type, forming micelles with core sizes ranging between 50 to 130 nm.^35^ Both studies found that the composite nanostructure produces stronger fluorescence signals than individual quantum dots. The enhancement in optical properties is likely due to surface modification that prevents oxidation and exciton dissociation, thus promoting radiative decay pathways.^36^ However, the use of hydrophobic components and heavy metals, such as cadmium in this case, is not preferred for *in vivo* bioimaging applications due to toxicity and clearance issues. In comparison with previous work, the platform described herein is entirely composed of biocompatible, non-toxic, hydrophilic materials that are hydrolytically degradable and excretable, thus representing a practical bioimaging agent with great translational potential. Furthermore, some of the photons absorbed can be converted into thermal energy (or ultrasound via thermoelastic expansion) through non-radiative decay, a process that is partly dependent on the nanoparticle structure.^37^ Previous reports have indicated that an increase in nanoparticle size, such as the clustering of small core gold nanoparticles in PCPP, would result in an aggregation induced red-shift toward higher wavelengths in the absorption onset.^12, 38^ Close packing of nanoparticles, such as ICG-gold nanoparticle clusters, may also improve photothermal conversion efficiency and thus increase non-radiative emissions, leading to greater output of heat and sound energy.^8, 19, 39^ Although the photothermal conversion efficiency of gold nanostructures is typically among the best of all PTT nanoprobes, the high material cost of gold is not ideal for widespread clinical use. In addition, gold nanorods are most explored for cancer PTT since their resonance peaks can be shifted toward the NIR spectrum,^40^ where tissue is more transparent to light; however, their synthesis usually involves the use of organic solvents and highly cytotoxic surfactants, such as CTAB.^41^ In comparison, our platform is much lower in cost with the use of a silver-based agent, and the improvement in photoconversion efficiency via polymeric encapsulation has resulted in relatively comparable therapeutic effects. Our platform is based on the use of aqueous solvents and non-toxic reagents, which eliminates the need for further surface modifications and ensuring safety in biomedical applications.

In our design of AgPCPP, we have loaded hydrophilic nanoparticles in PCPP-based hydrogel assemblies. PCPP nanoparticles can be loaded with other payloads in place of or in combination with Ag_2_S-NP to broaden their applicability. Such payloads may include hydrophilic drugs for synergistic chemotherapy,^42^ nanoparticles of high Z element for multi-energy CT imaging,^43^ or gold nanoparticles for radiotherapy.^1^ We can synthesize Ag_2_S-NP with fluorescence emission in the second NIR window (by increasing the core size) for imaging of deep tissues without significant effects from tissue absorption and background autofluorescence. Moreover, photothermal ablation has been found to activate sustained antitumor immune responses and produce *in situ* cancer vaccine-like effects that eliminate primary tumors and prevent further metastasis.^44^ Recently, multiple PTT cycles with Ag_2_S-NP have been shown to achieve greater immunotherapeutic effects compared to a single PTT-induced immune response.^45^ This type of photo-immunotherapy can potentially be performed using our agent since these nanoparticles are stable under several irradiation/cooling cycles, and their photoconversion activities are significantly improved with polymer encapsulation (*i.e.*, greater temperature elevation with less dose).

There are certain limitations to this study. For example, the safety of our nanoparticle formulations studied herein is limited to the cell lines and mouse models used. Further assessment of their biocompatibility besides cell viability assay (*e.g.*, DNA damage and free radical generation) and more extensive *in vivo* toxicology studies (*e.g.*, repeat dosing) would be desired. For future work, synthetic parameters (*e.g.*, anionic surface ligands) of Ag_2_S-NP will be closely studied to formulate AgPCPP with high payload yield in order to decrease the polymer used in synthesis and thus reduce the viscosity of the solution suitable for eventual human subject injection. *In vivo* intraoperative, image-guided surgical resection will be performed to determine the utility of our nanoplatform in that application, which was indicated by the *in vitro* phantom results. Longer imaging and biodistribution time periods (*e.g.*, 6-12 months) will be done to investigate the effect of degradability on contrast generation and the amount of time needed to achieve complete or sufficient clearance.

## Conclusion

In summary, we have developed a unique, multifunctional “one-for-all” contrast agent based on polymer encapsulated small, NIR fluorescent Ag_2_S-NP. For the first time, we have demonstrated the use of PEGylated PCPP copolymer in modulating the size of PCPP nanoparticles and for encapsulating contrast payloads. We have demonstrated good synthetic control of AgPCPP, efficient loading of hydrophilic payloads, and desired degradation rate *in vitro*. We have shown that the AgPCPP formulation is a robust contrast agent for X-ray and optical-based imaging modalities both *in vitro* and *in vivo*. AgPCPP are also advantageous over individual Ag_2_S-NP because AgPCPP produce greater photothermal heating due to increased NIR absorbance. Tumor growth was significantly inhibited by AgPCPP under NIR laser irradiation. Therefore, our promising results suggest that AgPCPP represent a biocompatible and biodegradable theranostic platform with potential translational impact.

## Materials and methods

### Materials

Silver nitrate (AgNO_3_, 99%), 2-mercaptopropionic acid (2-MPA, 95%), poly[di(carboxyphenoxy)phosphazene] disodium salt (PCPP, 1 MDa), spermine tetrahydrochloride, calcium chloride dihydrate (CaCl_2_), and sodium hydroxide solution (NaOH, 1 N) were purchased from Sigma-Aldrich (St. Louis, MO). Anhydrous sodium sulfide (Na_2_S) was purchased from Alfa Aesar (Tewksbury, MA). Herringbone microfluidic chip mixers and male luer fluid connectors were purchased from Microfluidic ChipShop (Jena, Germany). PEGylated derivative of PCPP, which contained 1% (mol) of methoxypolyethylene glycol amino (MW 5kDa) and 99% (mol) of carboxylatophenoxy side groups, was synthesized using previously described methods.^46^

### NIR fluorescent Ag_2_S-NP synthesis

The method to synthesize Ag_2_S-NP was adapted from a previous report.^19^ Briefly, 218 µL of 2-MPA (2.5 mmol) was added to a 250 mL conical flask containing 75 mL of DI water. The pH of the solution was adjusted to 7.5 using NaOH. Then, 42.5 mg of AgNO_3_ (0.25 mmol) was added to the flask, and the pH of the resulting solution was again adjusted to 7.5 using NaOH. A solution containing 5 mg of Na_2_S (0.0625 mmol) in 25 mL of DI water was prepared separately and slowly added to the above solution in a dropwise fashion. The reaction vessel was covered in aluminum foil and stirred for 24 hours under ambient conditions. The nanoparticle suspension, which appeared to be dark brown, was washed three times in DI water and concentrated to a final volume of 1 mL using 3 kDa MWCO ultrafiltration tubes (Sartorius Stedim Biotech, Germany). The tubes were spun at 4000 rpm for 40 minutes. Finally, the nanoparticles were filtered through a 0.02 µm membrane to remove any aggregates and stored at 4°C.

### AgPCPP nanoparticle synthesis

The procedure to synthesize PCPP nanoparticles formulated with PEG-PCPP copolymers was adapted from previous studies.^12, 13^ An example synthesis is as follows. A 2 mL solution containing PCPP (1.8 mg) and PEG-PCPP (0.2 mg) was prepared in PBS and loaded in a 10 mL syringe. A 2 mL solution containing spermine (1.96 mg) was also prepared in PBS and adjusted to pH 7.4 before loading in a separate 10 mL syringe. Both solutions were injected into a herringbone microfluidic chip mixer at a flow rate of 6 mL/min. The resultant mixture was quickly transferred to 100 mL of 8.8% (w/v) CaCl_2_ solution and stirred for 20 minutes. The solution was washed three times by centrifugation at 800 rcf for 10 minutes in DI water, and then stored at 4 °C in DI water for subsequent use. To form AgPCPP nanoparticles, the desired amount of Ag_2_S-NP was added to the PCPP and PEG-PCPP solution, while keeping the total volume fixed at 2 mL. The amounts of PEG-PCPP added in the formulation were varied from 0 to 0.5 mg to synthesize AgPCPP of varying sizes.

### Nanoparticle characterization

The core sizes and morphologies of Ag_2_S-NP and AgPCPP nanoparticles were examined using a Tecnai T12 transmission electron microscope (FEI, Hillsboro, OR) operating at 100 kV. The absorption spectra of the nanoparticles were characterized using a Genesys UV/visible spectrophotometer (Thermo Scientific, USA) after diluting with DI water. The NIR fluorescence emission spectra were recorded using a SpectraMax M5 microplate reader (Molecular Devices, San Jose, CA) with an excitation wavelength of 700 nm. The surface morphology and elemental content of AgPCPP nanoparticles were assessed using an FEI Quanta 600 scanning electron microscope (FEI, Hillsboro, OR) operated at 15 kV and equipped with EDAX EDS detectors (Ametek, Mahwah, NJ). The elemental silver concentrations were measured by ICP-OES (Spectro Genesis, Germany) following digestion in 1 mL of nitric acid and dilution with DI water. The infrared spectra of Ag_2_S-NP, AgPCPP, 2-MPA, PCPP, and PEG-PCPP were collected using a JASCO FT/IR-480 Plus spectrophotometer. The X-ray diffraction patterns of AgPCPP and Ag_2_S-NP were recorded using a Rigaku MiniFlex X-ray diffractometer operated at 45 kV, 30 mA, 1.5406 Å Cu Kα radiation wavelength, 2° per minute scan rate, and 20°-60° scan range.

### *In vitro* biocompatibility

The MTS assay (Promega, Madison, WI) was used to assess the cytotoxicity of Ag_2_S-NP and AgPCPP nanoparticles (*i.e.*, 10% of PEG-PCPP and 90% of PCPP (w/w)) towards the SVEC4 (endothelial cells), MDA-MB-231 (breast cancer cells), HepG2 (hepatocytes) and Renca (epithelial kidney cells). The cells were cultured according to the supplier recommendations. In accordance with a previously published method,^10^ the cells were seeded in a 96 well plate at a cell density of 10,000 cells per well and allowed to stabilize for 24 hours. Then, the cells were treated with the nanoparticles at different concentrations for 24 hours: 0 (control), 0.05, 0.1, 0.25 and 0.5 mg Ag per mL (*n* = 3 wells per plate used). Once the treatment was completed, the cells were washed with PBS twice and incubated with the assay, after which the absorbance at 490 nm was recorded. The relative cell viability (%) for each concentration and cell type was presented as mean ± SD. Experiments were done in triplicate.

### *In vitro* photothermal effects

Samples (1 mL) of ultrasmall Ag_2_S-NP and AgPCPP nanoparticles (10% PEG-PCPP) at a concentration of 0.5 mg Ag per mL were irradiated using an 808 nm laser (OEM Laser Systems, Midvale, UT) at a power density of 2 W per cm^2^ for 10 minutes in a quartz cuvette. Then, the samples were allowed to cool for 10 minutes without irradiation. The temperature was monitored throughout the entire experiment using an optical thermometer probe (Qualitrol Corporation, Fairport, NY). The photostability of AgPCPP nanoparticles was assessed by irradiating four times in succession using the same method. Additionally, AgPCPP nanoparticles were subjected to laser irradiation at different silver concentrations (0.125, 0.25 and 0.5 mg per mL) and laser power densities (0.5, 1.0, 1.5, 2 and 2.5 W per cm^2^) for 10 minutes during which changes in temperature were recorded. DI water was used as control.

A protocol from a previous study was used to evaluate cytotoxicity from photothermal ablation.^39^ MDA-MB-231 cells were seeded in a 96 well plate at a cell density of 10,000 cells per well with two empty wells between them. The following day, media was replaced with fresh 37°C media containing free Ag_2_S-NP or AgPCPP at various concentrations (*n* = 3 wells per condition). The wells were immediately irradiated with an 808 nm laser at a power of 1.5 W for a total of 5 minutes. The plates were then incubated for 24 hours, after which they were washed twice with PBS and analyzed using the MTS assay. The absorbance at 490 nm was recorded to determine the cell viability relative to control (%) at different conditions, which was presented as mean ± SD. Experiments were performed in triplicates.

### *In vitro* phantom imaging

Photoacoustic images were obtained using a Vevo LAZR system (VisualSonics Inc., Toronto, Canada) with an excitation wavelength of 700 nm. Ag_2_S-NP and AgPCPP nanoparticles (*n* = 3 per sample) were loaded into polyethylene tubing (40 µL) at a range of concentrations (0 to 1 mg Ag per mL). The tubes were placed in a plastic holder, which was immersed in DI water. The following parameters were used during image acquisition: PA gain 24 dB, ultrasound gain +27 dB, distance 1 cm from the LZ250 transducer (75 µm axial resolution; 13-24 MHz broadband frequency). The images were analyzed using ImageJ and the ROI values were recorded and normalized with respect to DI water. Data was presented as mean ± SD.

NIR fluorescence images were acquired using an IVIS Spectrum instrument (PerkinElmer, Waltham, MA) with excitation and emission wavelengths at 710 nm and 820 nm, respectively. Ag_2_S-NP and AgPCPP nanoparticles (*n* = 3 per sample) suspended in DI water at different concentrations (ranging from 0 to 5 mg Ag per mL) were placed in a black bottom 96 well plate (100 µL per well). The images were analyzed using Living Image 4.5.4 Software and the ROI values were recorded and presented as mean ± SD.

Intraoperative NIR fluorescence images were acquired using a dual-NIR channel Vet-FLARE imaging system (Curadel, Marlborough, MA). The system was equipped with a Model VP1 laser source and a Model VC2 handheld imaging head. Ag_2_S-NP and AgPCPP nanoparticles (*n* = 3 per sample) of varying concentrations (0 to 8 mg Ag per mL) were suspended in DI water and were loaded in 0.2 mL PCR tubes. The tubes were then placed in the center of the FLARE LT-1 light-tight enclosure. The following parameters were used during live mode image acquisition: power of 100%, exposure of 47 ms, camera gain of 1 v/v, mean brightness of -26, mean contrast of 20, mean gamma of 1. The color images, 2 independent NIR fluorescence images (700 nm and 800 nm) and merged images were acquired simultaneously. The 700 nm and 800 nm NIR fluorescence images were pseudo-colored blue and green, respectively, in merged images. ROI values were recorded using custom software and presented as mean ± SD.

CT imaging was performed using a MILabs µCT scanner (Utrecht, The Netherlands) with a tube voltage of 50 kV, tube current of 240 µA, step angle of 0.75°, and exposure time of 75 ms. Ag_2_S-NP and AgPCPP nanoparticles (*n* = 3 per concentration) as well as other control materials of varying concentrations (0 to 4 mg Ag per mL) were suspended in 1% agar gel and were loaded in 0.2 mL PCR tubes. The samples were secured in a plastic rack with parafilm. The identical samples were loaded in an acrylonitrile butadiene styrene tube holder and placed in the borehole of an anthropomorphic thorax phantom body (QRM GmbH, Mohrendorf, Germany) for imaging with a SOMATOM Force clinical CT scanner (Siemens Heathineers, Erlangen, Germany). The clinical CT images were acquired using an adult abdomen routine imaging protocol at 80, 100, 120, and 140 kVp. The following parameters were used during image acquisition: FOV = 37 × 37 cm, tube current = 360 mA, anode angle = 8°, exposure time = 0.5 s, slice thickness = 0.5 mm and matrix size = 512 × 512. For both µCT and clinical CT imaging, Osirix 64-bit software was used to measure the CT attenuation for each sample at each concentration and to calculate the attenuation rates. Data was presented as mean ± SD.

### AgPCPP degradation

AgPCPP nanoparticles (10% of PEG-PCPP and 90% of PCPP (w/w)) with a silver content of 0.1 mg were added to 1 mL of PBS with 10% FBS in micro-centrifuge tubes. The tubes were incubated in a water bath at 37 °C over a period of 7 days. At the desired time points, samples were centrifuged at 800 rcf for 10 minutes, and the supernatants and pellets were collected and analyzed via ICP-OES to measure the release of small Ag_2_S-NP. Three samples were prepared for each time point. The amount of released Ag_2_S-NP was calculated by dividing the mass of silver measured in the supernatant by the total silver mass measured in both the supernatant and the pellet. The pellet from a sample incubated for 7 days was examined under TEM and SEM to confirm the degradation of AgPCPP nanoparticles.

### Animal and tumor model

All animal studies were conducted with approval by the Institutional Animal Care and Use Committee of the University of Pennsylvania under protocol number 807141. Female nude mice, aged approximately 6-8 weeks, were obtained from Jackson Laboratory (Bar Harbor, ME). These mice were inoculated with MDA-MB-231 breast cancer cells (1 × 10^6^ cells per mouse, 50 µL total volume in HBSS) by injecting orthotopically into the fourth abdominal mammary pad above the right flank only. Four mice were used per group (n = 4) in each experiment.

### *In vivo* multimodality imaging

All tumors were grown to an approximate volume of 100 mm^3^ and pre-contrast tumor images were acquired at this time. Nanoparticles (2 mg Ag per ml, 30 µL total volume) were injected directly into the tumors. The mice were scanned before injection and at 1, 72, and 168 hours post injection. Mice were anesthetized with isoflurane during all imaging experiments.

CT images were acquired with a MILabs µCT scanner (Utrecht, The Netherlands) using the following parameters: kVp = 50 kV, current = 240 µA, step angle = 0.75°, and exposure time = 75 ms. The images were analyzed using OsiriX. An ROI was placed over the tumor site of 2D axial plane images, and CT attenuation values from three slices were recorded. CT attenuation of tumors at each time point is presented as mean ± SEM in Hounsfield units.

Photoacoustic images were acquired with a VisualSonics Vevo Lazr (Toronto, Canada) using the LZ250 transducer. Bubble-free gel was applied in between the transducer and tumor surface. The following acquisition parameters were used: PA gain 30-40 dB, priority 95%, and distance 12 mm. An ROI was placed on the photoacoustic signal of tumor site and the intensity was recorded over a range of wavelengths. Pre-injection scan, designated as background, was used as control.

NIR fluorescence images were acquired using an IVIS Spectrum instrument (PerkinElmer, Waltham, MA) with an excitation wavelength of 745 nm and an emission wavelength of 820 nm. An exposure time of 0.5 second was chosen. All images had the same illumination settings and were analyzed using the LivingImage software. The average radiant efficiency of the tumor area was recorded using the ROI tool and presented as mean ± SEM.

### *In vivo* photothermal treatment

When the tumor size reached 50 mm^3^ in volume, mice were anesthetized with isoflurane and nanoparticles (2 mg Ag per ml, 30 µL total volume) were injected directly into the tumors. Control groups included mice that were injected with saline and those that were not exposed to laser. Their body temperature was maintained using a temperature-controlled heating pad. Immediately after injection, tumors were subjected to photothermal ablation using an 808 nm laser for 30 minutes at a power density of 0.7 W per cm^2^ (1.2 cm in laser diameter or 1.13 cm^2^ laser area, which fully covered all tumor surface). A FLIR ONE (Wilsonville, OR) thermal imaging camera was used to monitor tumor and chest surface temperatures before and during laser irradiation. Infrared thermography acquired at various time points was analyzed for maximum tumor temperature as well as tumor temperature normalized to overall body temperature. Tumor volume and animal weight were measured in all groups and recorded for 14 days after photothermal treatment. Animals with tumor length surpassing 15 mm at any given time were considered non-surviving. Animals were monitored at least daily for signs of distress and changes in eating pattern and behavior after photothermal treatment. Data are presented as mean ± SEM.

### Histopathology

Mice were injected with PBS solution or AgPCPP nanoparticles at a dose of 25 mg Ag per kg and were sacrificed after 24 hours and 7 days. Their major organs were collected, rinsed with cold PBS, and cut into 5 mm thick pieces. The organ pieces were fixed in 10% buffered formalin overnight at 4 °C. The samples were processed and stained with hematoxylin and eosin (H&E) by the Children’s Hospital of Philadelphia Pathology Core.

### Serum biochemistry

Whole blood samples (200 µL total volume) were collected from mice via cardiac puncture. The blood was allowed to clot by leaving it undisturbed at room temperature for 30 minutes. The clot was then removed by centrifugation (1000 rcf for 10 minutes), and the resulting supernatant or serum was collected and stored at -20 °C until further analysis. Serum biomarkers were analyzed by IDEXX Bioanalytics (North Grafton, MA). Data are presented as mean ± SEM.

### Biodistribution

A subset of tumor-bearing mice was injected with Ag_2_S-NP or AgPCPP nanoparticles (25 mg Ag per kg, 190 µL total volume) via the tail vein. At 24 hours post injection, mice were euthanized and blood samples were collected from them, after which the organs were perfused with 20 ml of PBS. Major organs including the heart, lungs, liver, kidneys, spleen, bladder, tumor, and feces were collected and weighed. The tissues were minced and digested in nitric acid overnight. The samples were then diluted with DI water and filtered before measuring the concentration of silver with ICP-OES. For longer term biodistribution analysis, two sets of non-tumor-bearing mice were injected with AgPCPP nanoparticles using the same conditions as above. Their organs were collected at 1 day and 7 days post-injection and analyzed using the aforementioned method. Data are presented as mean ± SEM.

### Examination of liver tissue via TEM

Liver tissues from mice treated with AgPCPP nanoparticles (25 mg Ag per kg) were fixed in 2.5% glutaraldehyde and 2% paraformaldehyde, after which were stained and embedded. The sample sections were mounted onto copper grids, which were viewed using a Tecnai T12 electron microscope operated at 100 kV.

### Statistical analysis

All studies were conducted in triplicate or in three independent experiments. GraphPad Prism 8 software was used to perform all statistical analyses. The figure caption contains the specific statistical tests that were performed, when applicable.

## Acknowledgements

This material is based upon work supported by the Brody Family Medical Trust Fund Fellowship (JCH). Partial support was provided by the NIH under grants R01CA227142 (DPC), R25CA140116 (UPenn SUPERS program), and K99EB028838 (MH). In addition, this work was carried out in part at the Singh Center for Nanotechnology, which is supported by the NSF National Nanotechnology Coordinated Infrastructure Program under grant NNCI-2025608. The authors thank the Electron Microscopy Resource Laboratory at the University of Pennsylvania, especially Sudheer Molugu and Biao Zuo for their support with TEM. We also thank the Small Animal Imaging Facility at the University of Pennsylvania, including Eric Blankemeyer for his help with micro-CT imaging, Ching Huang for her help with optical imaging, Susan Schultz for her help with PA imaging, and Joann Miller for her help with PTT experiments.

## Notes

### Competing Interest Statement

The authors have declared no competing interest.

