## Supplementary Information for "A biodegradable “one-for-all” nanoparticle for multimodality imaging and enhanced photothermal treatment of breast cancer"

*Corresponding author

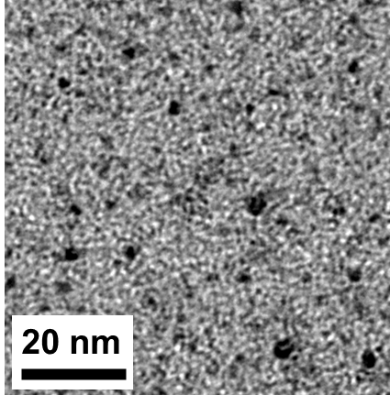

**Figure S1.** TEM image of small core Ag_2_S-NP.

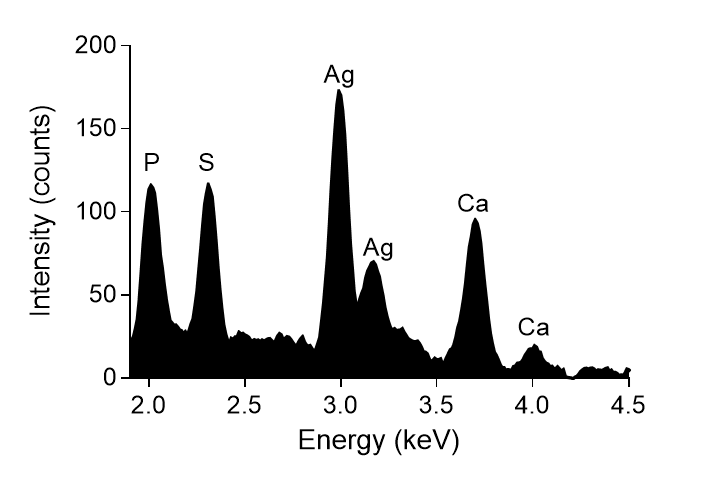

**Figure S2.** EDX elemental spectrum of AgPCPP nanoparticles.

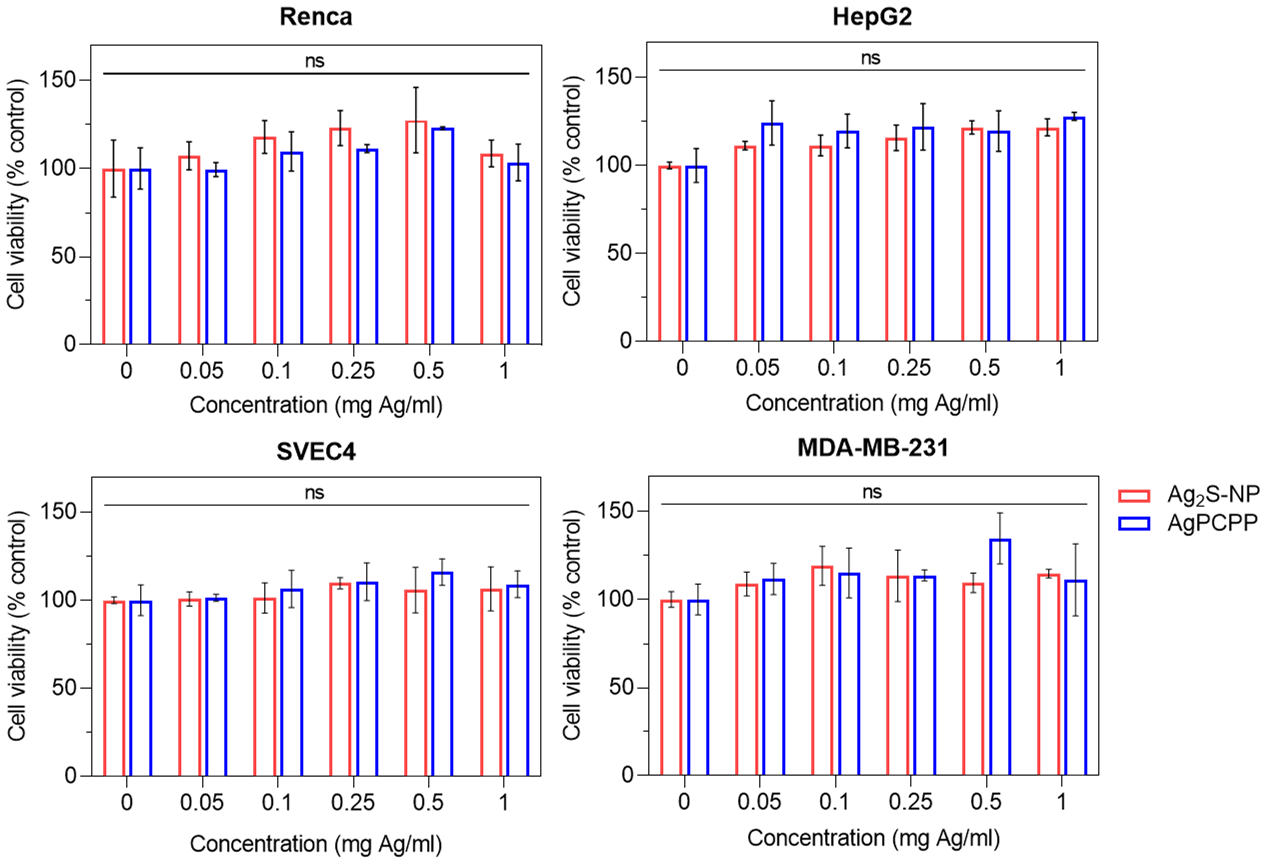

**Figure S3.** *In vitro* cytocompatibility of Ag_2_S-NP and AgPCPP nanoparticles. The four cell lines used for safety assessment were Renca (kidney epithelial cells), HepG2 (hepatocytes), SVEC4 (endothelial cells), and MDA-MB-231 (breast cancer cells). Data are presented as mean ± SD. Error bars are one SD. ns = not significant (one-way ANOVA with Tukey’s multiple comparisons test).

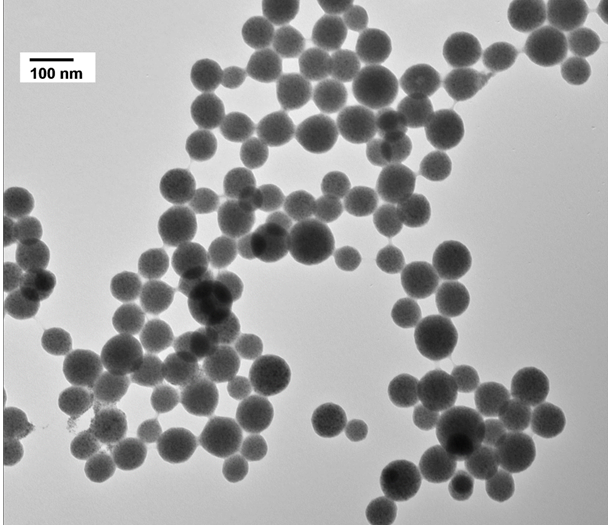

**Figure S4.** TEM of AgPCPP nanoparticles following repeated cycles of laser irradiation.

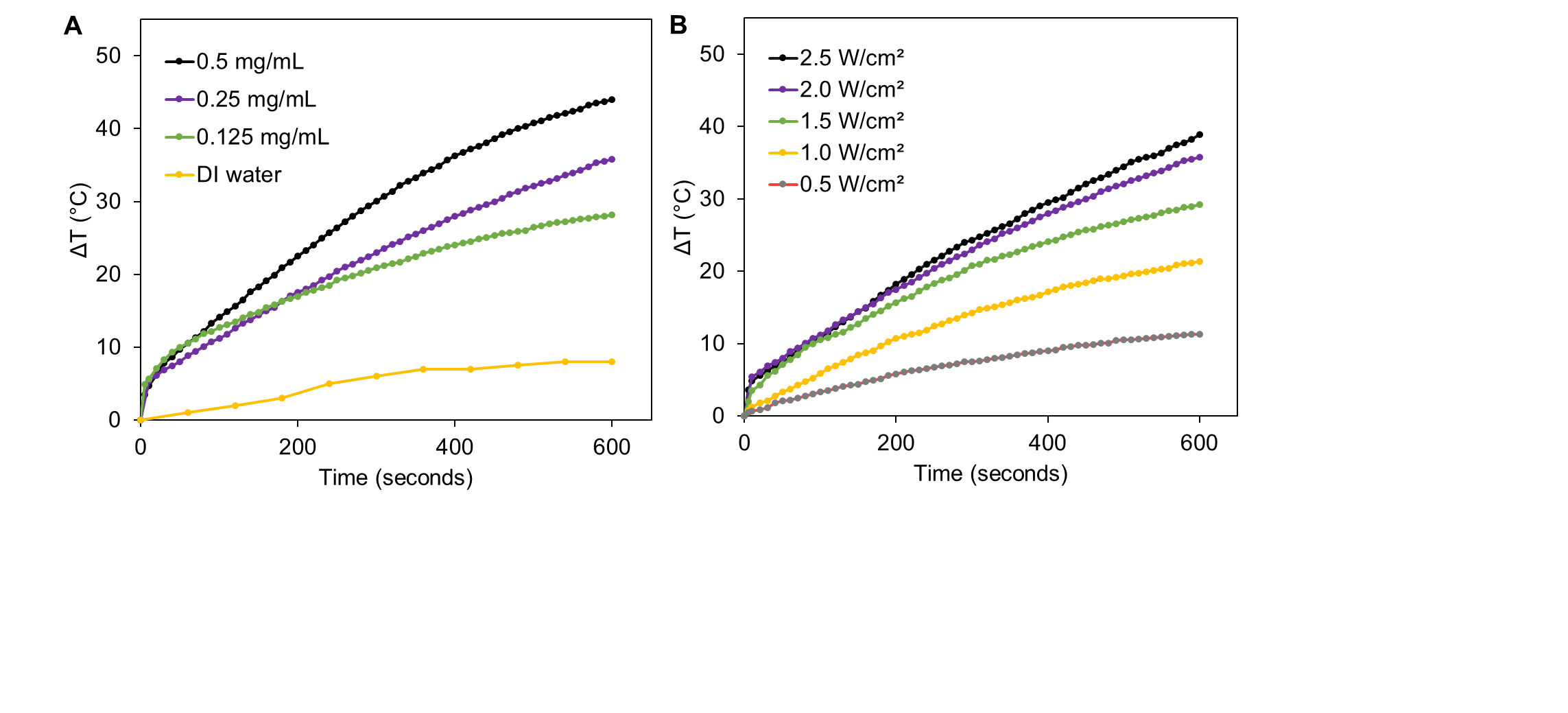
**Figure S5.** Temperature elevations by AgPCPP are dependent on (A) the concentration of silver in the solution and (B) laser power density (at a concentration of 0.25 mg Ag per ml).

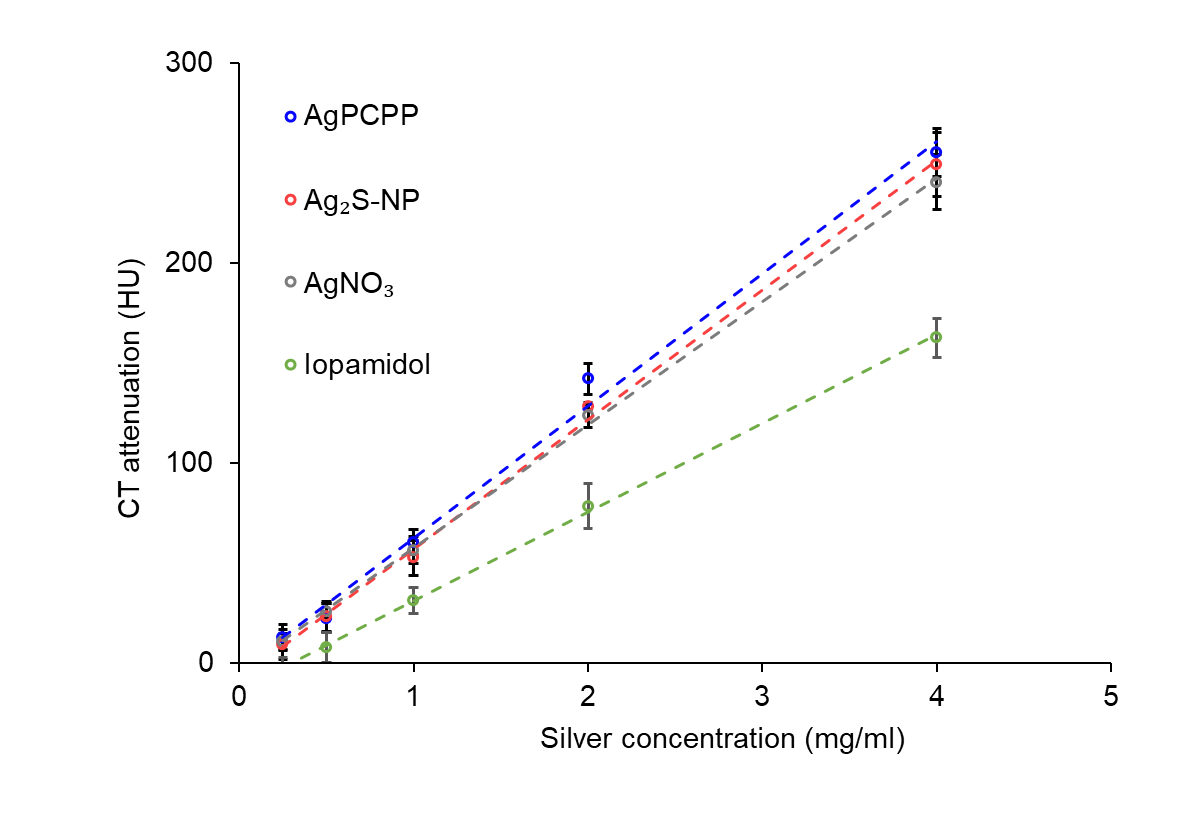

**Figure S6.** CT attenuation of various contrast materials as a function of concentration.

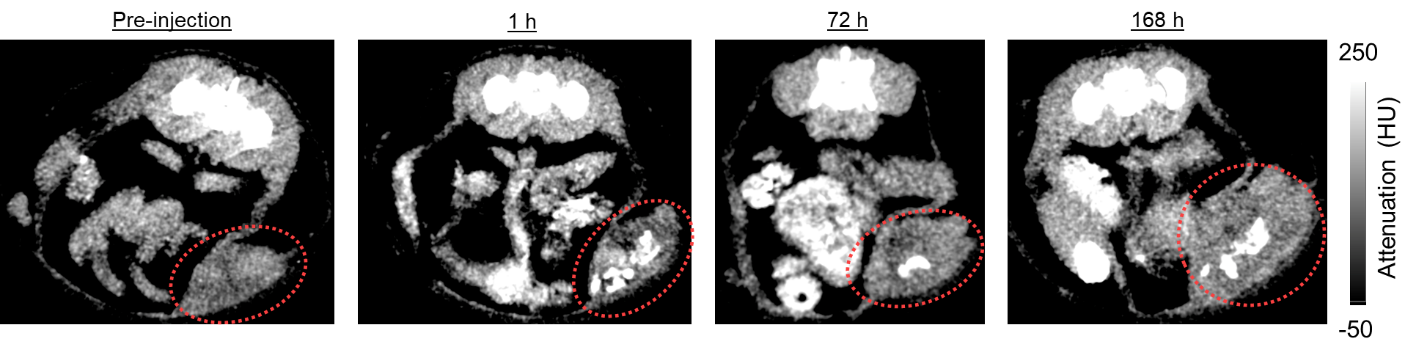
**Figure S7.** CT images of AgPCPP nanoparticles injected in the tumors viewed in axial plane. Red dashed circles indicate tumor site.

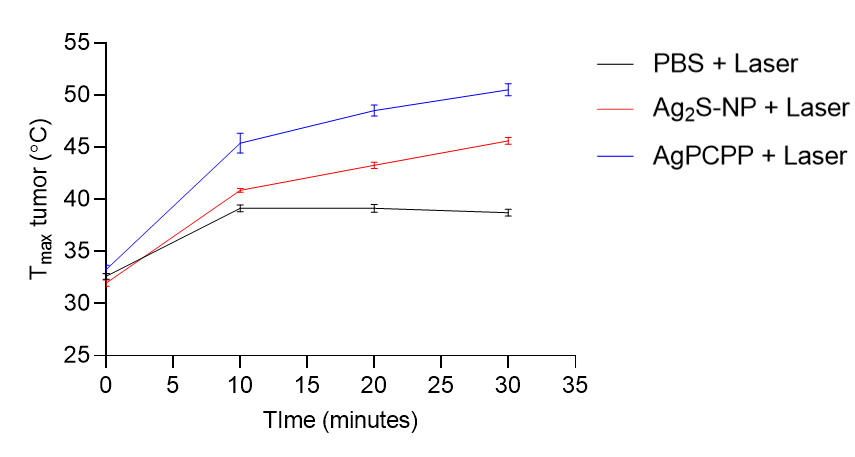

**Figure S8.** Peak tumor temperature achieved by PBS, Ag_2_S-NP, and AgPCPP nanoparticles under laser irradiation (mean ± SEM).

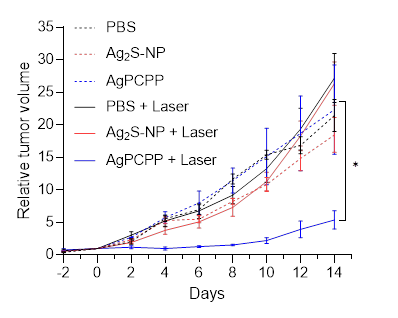

**Figure S9.** Tumor growth curves of mice treated with saline, Ag_2_S-NP, and AgPCPP nanoparticles with or without 30 minutes of laser irradiation for a total of 14 days post therapy (mean ± SEM). day 0 = tumor injection and laser treatment. **p* < 0.05 (two-way ANOVA with Bonferroni’s multiple comparisons test).

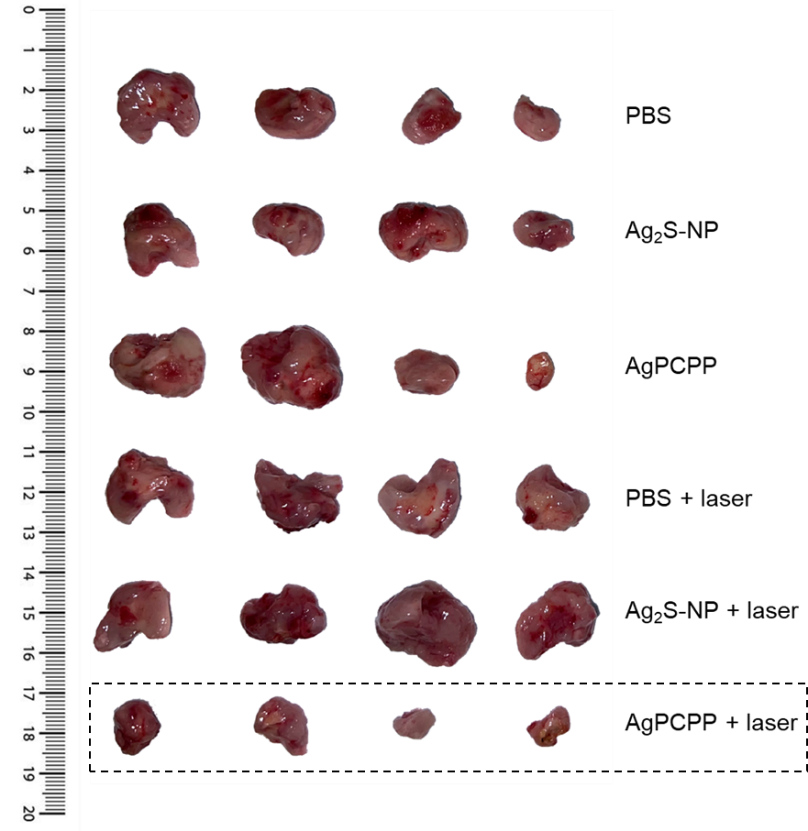

**Figure S10**. Photograph of tumors extracted from the mice at 14 days post treatment.

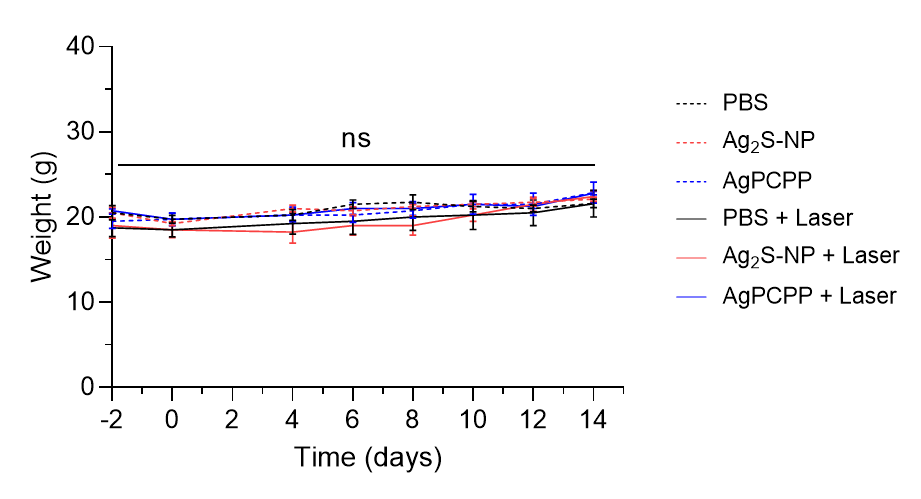

**Figure S11.** Body weights of mice before and after receiving treatments. Data are presented as mean ± SEM. ns = not significant (two-way ANOVA with Bonferroni’s multiple comparisons test).

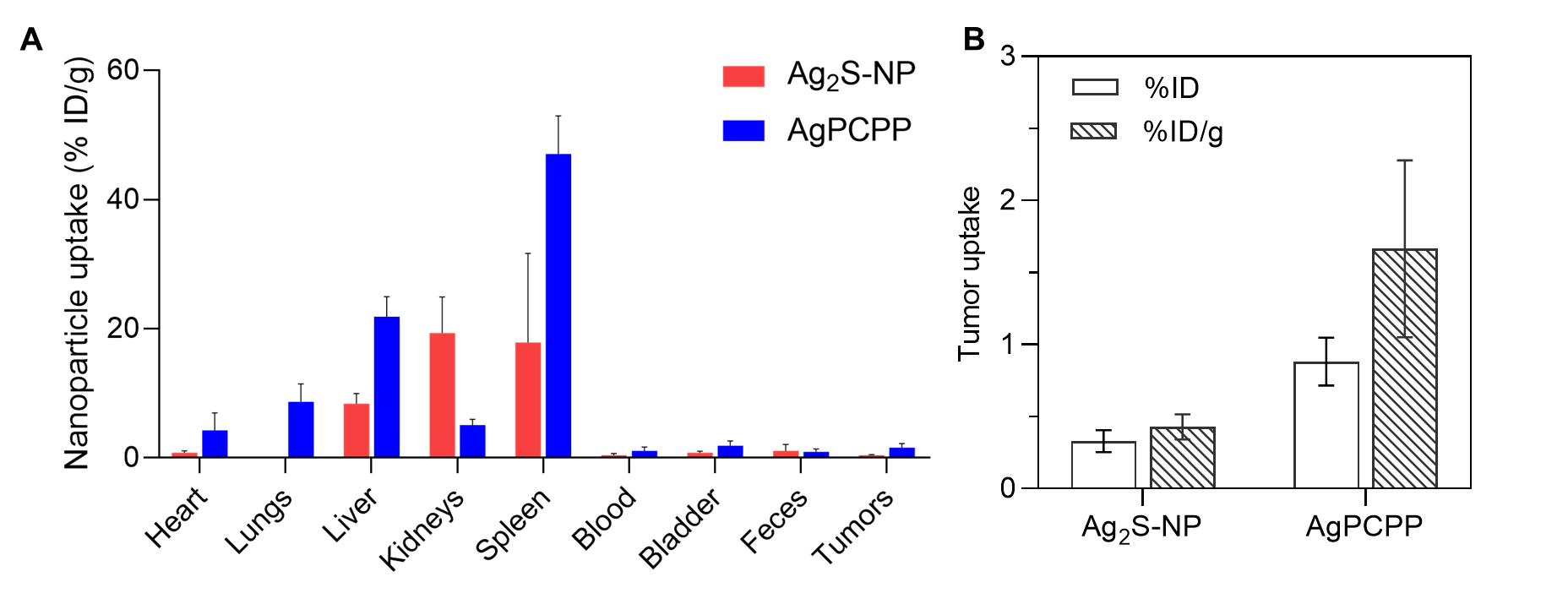

**Figure S12.** (A) Biodistribution profiles of Ag_2_S-NP and AgPCPP at 24 hours after intravenous administration. (B) Tumor uptake for both nanoparticle formulations. Data are presented as mean ± SEM.
